# Modelling rate-independent damping in insect exoskeleta via singular integral operators

**DOI:** 10.1101/2024.10.20.619287

**Authors:** Arion Pons

## Abstract

In insect locomotion, the transmission of energy from muscles to motion is a process within which there are many sources of dissipation. One significant but understudied source is the structural damping within the insect exoskeleton itself: the thorax and limbs. Experimental evidence suggests that exoskeletal damping shows frequency (or, rate) independence, but investigation into its nature and implications has been hampered by a lack methods for simulating the time-domain behaviour of this damping. Here, synergising and extending results across applied mathematics and seismic analysis, we provide these methods. We show that existing models of exoskeletal rate-independent damping are equivalent to an important singular integral in time: the Hilbert transform. However, these models are strongly noncausal, violating the directionality of time. We derive the unique causal analogue of these existing exoskeletal damping models, as well as an accessible approximation to them, as Hadamard finite-part integrals in time, and provide methods for simulating them. These methods are demonstrated on several current problems in insect biomechanics. Finally, we demonstrate, for the first time, that existing rate-independent damping models are not strictly dissipative: in certain circumstances they are capable of generating negative power without apparently storing energy, likely violating conservation of energy. This work resolves a key methodological impasse in the understanding of insect exoskeletal dynamics and offers new insights into the micro-structural origins of rate-independent damping as well as the directions required in order to resolve violations of causality and the conservation of energy in existing models.

## 1. Introduction

Exoskeletal structures play crucial roles in insect locomotion. In winged insects (Pterygota), the wings themselves are exoskeletal outgrowths originating from ancestral structures of uncertain form (Engel et al., 2013); and the thoracic exoskeleton contains not only the mechanisms that translate the action of the flight muscles into wingbeat motion (Melis et al., 2024; Walker et al., 2014); but also, in certain cases, structures that can store elastic energy and thereby reduce wingbeat power requirements (Gau et al., 2019; Warfvinge et al., 2017). In walking, jumping and hopping insects across orders, the exoskeleton of the legs and thorax provides contact and support forces, and can contribute to energy storage for use in jumps and bounds (Bolmin et al., 2021; Burrows, 2010). Studies of the energetics of insect locomotion focus often on the elastic properties of these exoskeletal structures—it is, after all, these elastic properties that enable improvements in flight efficiency (Gau et al., 2019; Pons et al., 2023) and jumping performance (Bolmin et al., 2021; Burrows, 2010). But damping is also at work: exoskeletal structures also dissipate energy via viscoelastic effects within the partially-sclerotised chitin-protein matrix of which they are composed (Aberle et al., 2017). Empirical evidence from dynamic mechanical analysis (DMA) across a range of species and locomotor systems—including hawkmoth flight motors (Gau et al., 2019; Wold et al., 2023); beetle elytra (Lomakin et al., 2010); and cockroach legs (Dudek and Full, 2006; Dudek and Full, 2007)— indicates that this damping is often largely frequency- or rate-independent: a property consistent with the characteristics of chitin and other cuticle polymers (Aberle et al., 2017; Martin and T., 1962; Sun et al., 2016), and which can lead to favourable control properties (Dudek and Full, 2007; Wold et al., 2023).

Linear rate-independent damping—otherwise termed structural or frequency-independent damping—permits a straightforward and convenient model representation in the frequency-domain: a constant imaginary value appended to the stiffness term within the force-displacement transfer function, often styled *iγ* (Dudek and Full, 2006; Gau et al., 2019; Wold et al., 2023). However, this straightforwardness conceals serious practical and analytical problems. Exact time-domain formulations of rate-independent damping are absent from biomechanical literature, and indeed, one finds this coefficient in no table of inverse Fourier transforms. Without such formulations, there is no accurate way to identify the classical damping parameter *γ* from time-domain exoskeletal mechanical data—particularly, when nonlinear effects are present, and frequency-domain analysis breaks down (Lynch et al., 2021; Pons and Beatus, 2022a). Neither is there any way to simulate time-domain models of the exoskeleton, *e*.*g*., as part of a full bio-aero-structural model of an insect flight motor (Gau et al., 2023; Pons, 2023).

In this work, we address all these issues by developing time-domain formulations of linear rate-independent damping, and applying them to problems in the identification of insect exoskeletal damping—a process based on methods across applied mathematics, seismic analysis, and biomechanics. First, in §2, leveraging techniques from seismic analysis (Inaudi and Makris, 1996; Makris, 2017), we develop an exact time-domain formulation for rate-independent damping based on the singular integral known as the Hilbert transform. In doing so, we determine this damping it to be non-causal—that is, in violation of the directionality of time. Following, and extending, further existing results (Makris, 1997a; Pons, 2024), we resolve the causality violation and obtain a pair of approximate causal time-domain formulations—which we identify, for the first time, as related non-smooth singular integrals. In §3 we develop new and highly accessible numerical methods for computing the responses of these singular-operator models of rate-independent damping, and for simulating models of the insect exoskeleton in which they are present.

Using these methods, in §4 we demonstrate a crucial and previously unknown caveat of linear rate-independent damping: such damping is not always dissipative, that is, it is capable of releasing energy, as well as absorbing it. In doing so we identify new restrictions on the use and validity of these damping models. Finally, in §5 we apply our techniques to challenging problems in entomological literature, including the identification of rate-independent damping parameters from exoskeletal transient responses and DMA data when structural nonlinearities and motion asymmetries are present (Dudek and Full, 2007; Wold et al., 2023); and time-domain simulation of structural models with rate-independent damping (Dudek and Full, 2007; Gau et al., 2023). In §6 we discuss these results and identify pathways toward more clearly understanding the mechanistic origins of and best-practice models for rate-independent damping. Synergising applied mathematics, seismic analysis, and insect biomechanics, our results provide wider integrative studies of insect locomotion with tools to integrate rate-independent damping within time-domain models of insect locomotion processes; as well as a more detailed understanding of the limits of validity of such models and the ways in which they can break down.

## 2. Formulations of rate-independent damping

### 2.1. Motivation and frequency-domain formulation

The challenges that arise in existing models of rate-independent damping in the insect exoskeleton are given context by the unique circumstances in which these models have been constructed. Consider a classical single degree-of-freedom (1DOF) viscous damper—for instance, as might initially be taken to model the structural damping associated with thoracic deformation in Dipteran and Hymenopteran flight (Fig. 1A) (Lynch et al., 2021; Wold et al., 2023). This damper can be expressed in the frequency (Ω ∈ ℝ) and time (*t* ∈ ℝ) domains as:

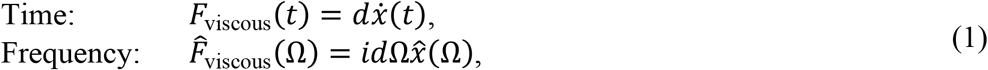

where *d* is the viscous damping parameter, and *F* is the action force—the force required to displace the system by displacement *x*. Note that the reaction force is −*F*, and that the relationship between time and frequency domains are given by the Fourier and inverse Fourier transforms—for generic function *G*(*t*):

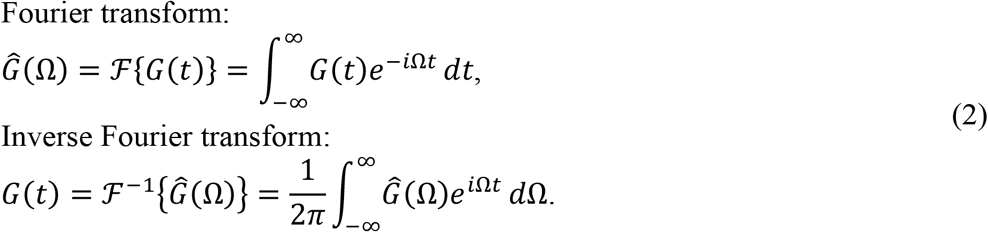

**Figure 1.**
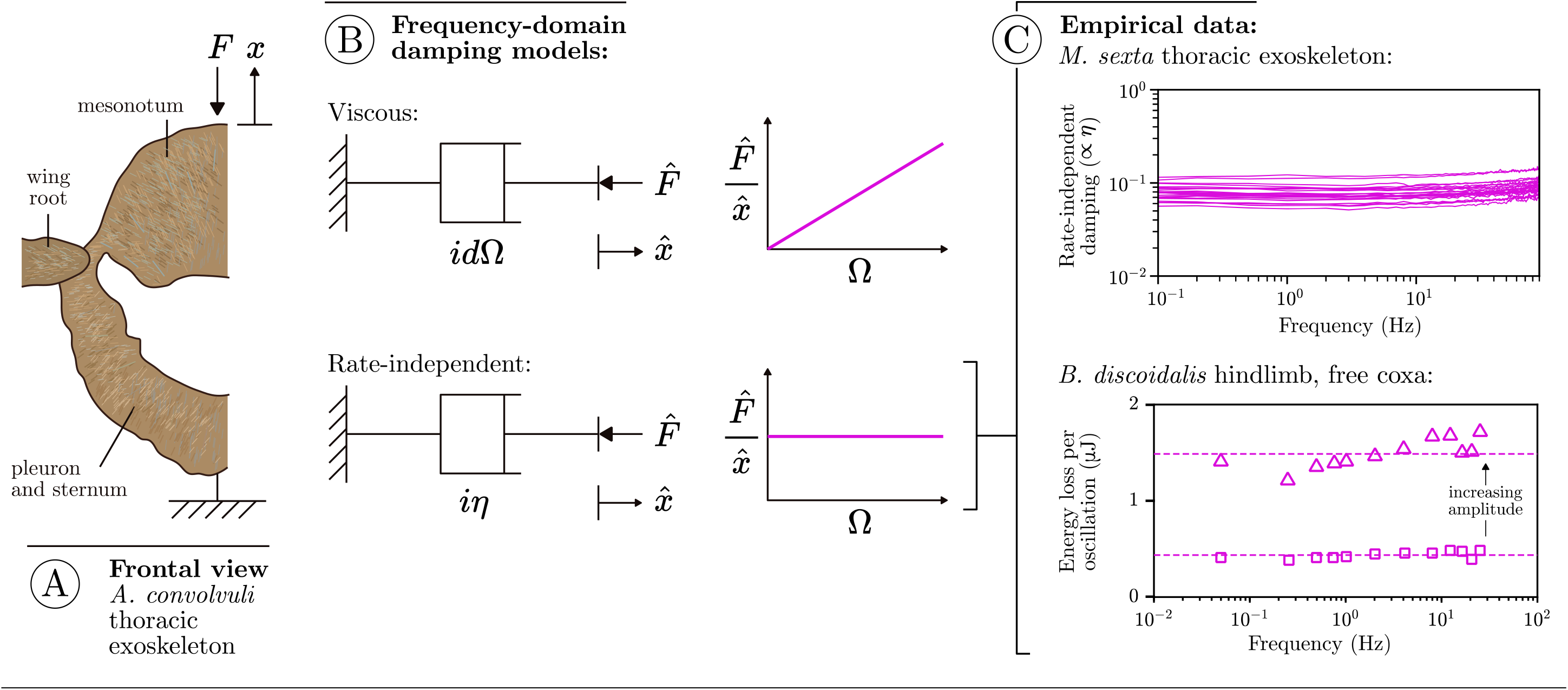
Context for rate-independent exoskeletal damping. (**A**) Frontal view of the thoracic exoskeleton of the hawkmoth *Agrius convolvuli*, after Ando et al. (2022), illustrating a DMA context for structural damping within hawkmoth thoraces during flight (Wold et al., 2023). (**B**) Viscous and rate-independent models of thoracic damping, with associated scaling of force with frequency (Ω). (**C**) Reported DMA data for several exoskeletal structures indicates rate-independent scaling: the thorax of the hawkmoth *Manduca sexta* (Gau et al., 2019); and the hindlimb of the deathhead cockroach *Blaberus discoidalis* (Dudek and Full, 2006).

In Eq. 1, the damping force is frequency-dependent: 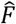 scales by Ω (Fig. 1B). Empirical data for a range of different insect exoskeletal structures contradicts this viscous model (Dudek and Full, 2006; Dudek and Full, 2007; Gau et al., 2019; Lomakin et al., 2010; Wold et al., 2023), and indicates rather than this force amplitude (and associated metrics, such as the energy loss per oscillation) is largely independent of frequency, over several orders of magnitude. Several such data are illustrated in Fig. 1C.

To capture rate-independent damping, common practice (Dudek and Full, 2006; Lynch et al., 2021; Pons, 2023) is to make an *ad hoc* modification to Eq. 1, deleting the factor of Ω:

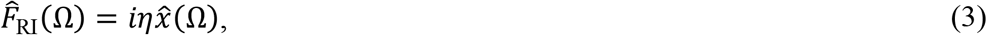

and defining a new rate-independent damping parameter *η* (Fig. 1B). This damping model is often combined with a linear stiffness 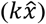 to form a Voight-type viscoelastic model, and in this case damping can be expressed as a dimensionless factor of stiffness (*η* = *kγ*). Estimates of *γ* range over 0.1-0.5 for different exoskeletal structures under different loading conditions (Dudek and Full, 2007; Wold et al., 2023). Formally, the modified model of Eq. 3 is well-defined for a single-sided Fourier series (positive Ω), but not for the complete double-sided Fourier transform (positive and negative Ω). For 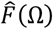 to inverse Fourier transform to a real-valued force *F*(*t*), the real part of 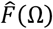 must be an even function; and the imaginary part an odd function. To ensure this while maintaining rate-independent damping, we must define an extension to Eq. 3:

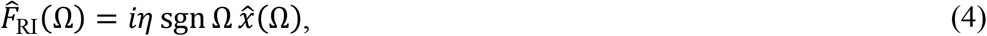

where sgn · is the signum function. Eq. 4 retains rate-independent force amplitudes, is equivalent to Eq. 3 for positive (‘physical’) frequencies, and is the formally-correct ideal rate-independent damping model over Ω ∈ ℝ, as used in seismic analysis (Inaudi and Makris, 1996; Luo and Ikago, 2021; Reggio and De Angelis, 2015).

### 2.2. The ideal rate-independent damper

The exact time-domain formulation of the rate-independent model, Eq. 4, is known in seismic analysis literature. To sketch the origins of this formulation, we observe that the Fourier transform is related the Hilbert transform, ℋ{·}, a real-valued singular integral transform representing the convolution (*) of a signal with the kernel 1/*πt* (Poularikas, 2018):

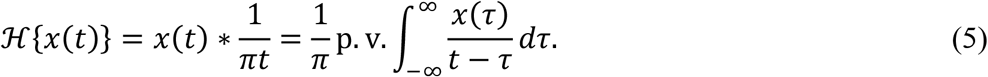

The kernel 1/*πt* is smooth and strongly singular, and so in general the value of Eq. 5 is determined via Cauchy principal value (p. v.). A relationship between Hilbert and Fourier transforms, in the function spaces where both are defined, can be expressed (King, 2009):

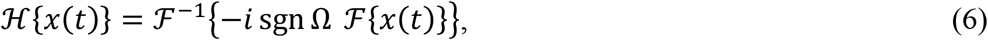

and so Eq. 6 can be rearranged into an exact and unique time-domain formulation for Eq. 4:

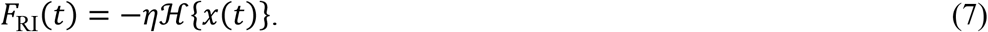

That is, the force response of the rate-independent damper is the Hilbert transform of the input displacement—as has been derived previously by seismic analysts (Inaudi and Kelly, 1995; Keivan et al., 2018). For illustration, Fig. 1 compares the response of a rate-independent damper (Eq. 7) to that of a viscous damper (Eq. 1), for a triangle wave input. A wide range of analytical results for the Hilbert transform have been tabulated (King, 2009; Poularikas, 2018), and numerical implementations are available in many scientific computing languages—we discuss these in more detail in §3.1. However, prior to this, two properties of the Hilbert transform should be noted:

i. The Hilbert transform term, −ℋ{·}, takes each harmonic component within the input, *x*(*t*), and shifts its phase by 90°. A simple-harmonic input thus leads to a simple-harmonic output: −ℋ{sin Ω*t*} = cos Ω*t* (Ω > 0), *i*.*e*., the output is identical to the derivative (Ω cos Ω*t*) but without the amplitude scaling by frequency: a model of rate-independent damping. However, for a general multiharmonic input, shifting harmonic components by 90° leads to a waveform that differs both from the original input and from the derivative. This feature will be important when we consider the damper’s energetics in §3.
ii. To evaluate the Hilbert transform at any instant a convolution must be performed over all time: past and future (Eq. 5). The ideal rate-independent damper is non-causal, *i*.*e*., it does not respect the directionality of time (Makris, 1997b). One mechanism to observe this non-causality is the damper’s memory function (Enelund and Olsson, 1999; Makris, 1997b), *Q*_RI_(*t*), representing its response to *δ*(*t*), a Dirac delta distribution at *t* = 0. As such, *Q*_RI_(*t*) is the inverse Fourier transform of the damper’s transfer function, and defines any time-domain response via convolution:

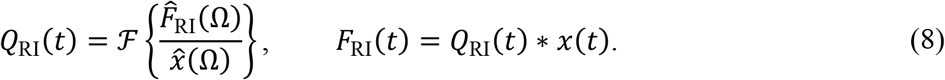

Hence:

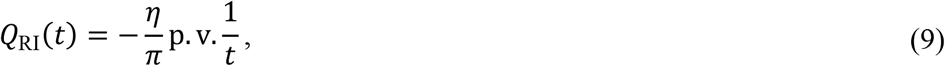

where p. v. 1/*t* is the generalised function defined by Cauchy principal value regularisation of the singular ordinary function 1/*t* (Kanwal, 2004; King, 2009). Eq. 9 is illustrated in Fig. 2: its significance is that it indicates that this damper is *strongly* non-causal: the damper weights information from the past and future equally. This non-causality is significant practical and physical hurdle: it makes time-domain simulation of initial value problems, such as flight motor models (Gau et al., 2023; Lynch et al., 2021), challenging; and by its nature we know that it cannot represent the behaviour of any physical structure.

**Figure 2.**
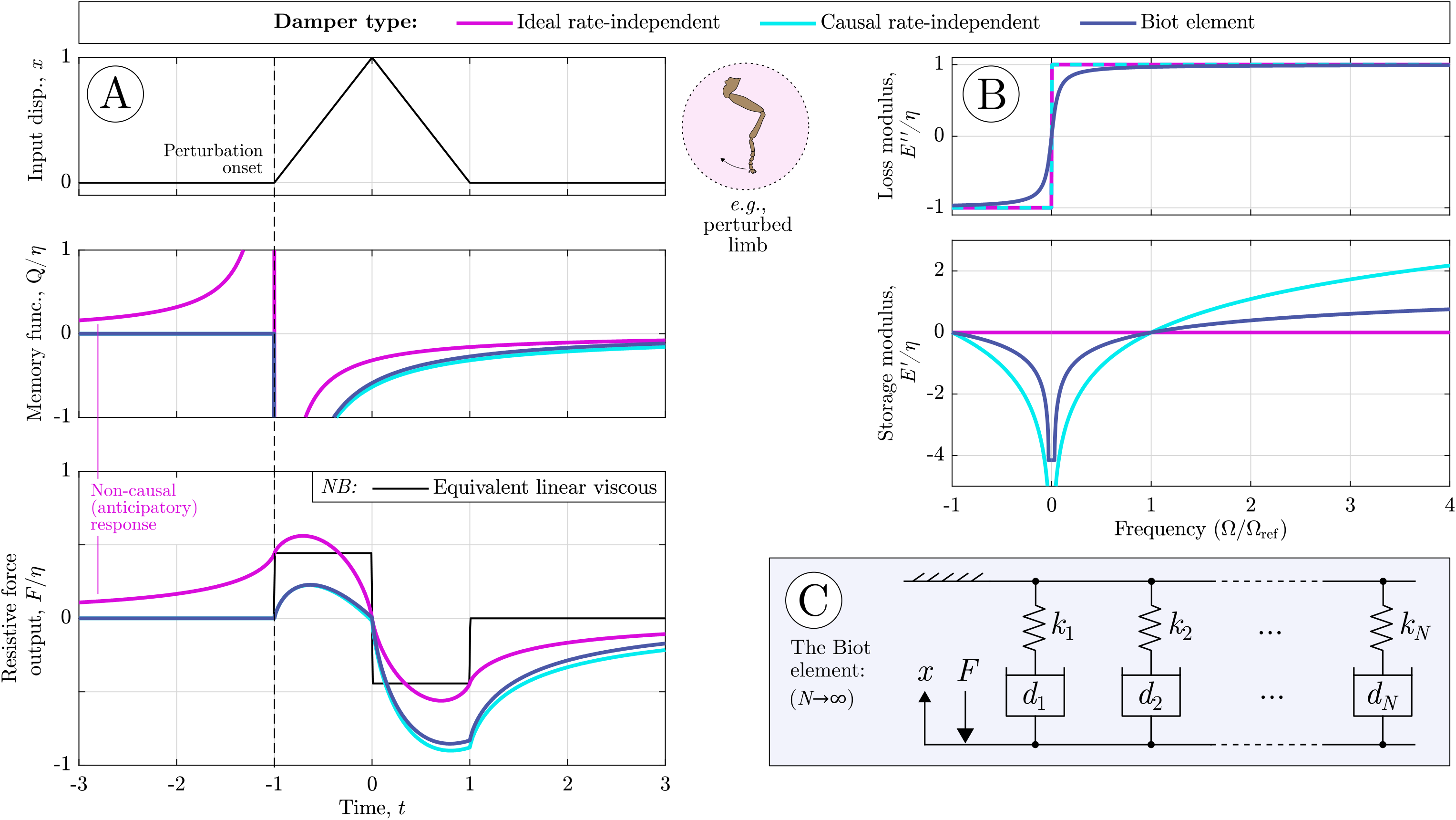
Behaviour of rate-independent dampers: ideal, causal, and the Biot element. (**A**) Time-domain damping responses to a triangle pulse, illustrative of a transient perturbation to an insect limb. Damper memory functions (*Q*/*η*, about perturbation onset) and force responses (*F*/*η*) are shown, computed via the numerical methods described in §3, and for tuning parameter values Ω_ref_ = π/2 and *N*_ref_ = 20 (see also §4.2). In these, the non-causal behaviour of the ideal rate-independent damper can be clearly seen. A linear damper of equivalent net power dissipation is included for comparison. (**B**) Loss and storage moduli of these dampers. Together, causality and loss modulus rate-independence strictly necessitate a logarithmic storage modulus. (**C**) Notably, the Biot element can be constructed as the infinite sum of linear stiffnesses and viscous dampers—illustrating a physical mechanism via which rate-independent damping can arise, as we discuss further in §6.1.

### 2.3. The causal rate-independent damper

It is natural then to consider modifying this model of ideal rate-independent damper to make it causal. As per classical viscoelastic analysis, any frequency-domain relationship between force 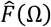 and displacement 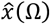 can be expressed via a complex modulus 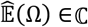, with the storage (*Ê*′ ∈ ℝ) and loss moduli (*Ê*″ ∈ ℝ) being the real and imaginary parts of this complex modulus:

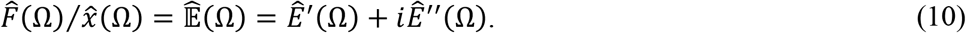

Physically,*Ê*′ loosely approximates structural stiffness, and *Ê*″ structural damping. But not every *Ê*′-*Ê*″ pair respects causality: rather, causal structures show a specific pair relationship, given by Titchmarsh’s theorem and its generalisations: loosely, these moduli must be Hilbert transforms (in Ω) of each other, with the caveat that certain free variables can arise, leading to a nonunique relationship (Nussenzveig, 1972; Pons, 2024).

The practical implication of causality analysis is that the rate-independent loss modulus 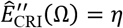 sgn Ω directly implies a certain storage modulus, 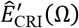. Previous analyses (Makris, 1997a; Pons, 2024) have derived this storage modulus as:

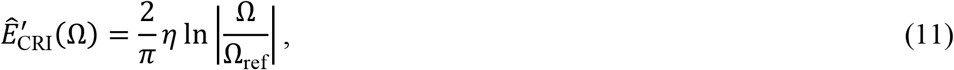

where Ω_ref_ > 0 is a free parameter—the reference frequency. Fig. 2 illustrates this causal storage modulus. The complete model of a causal rate-independent damper is thus:

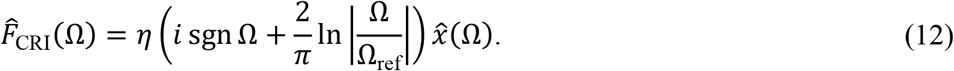

which is weakly singular at Ω = 0. We observe several properties of this model:

i. The reference frequency, Ω_ref_, functions as a linear stiffness parameter, setting the storage modulus to zero at Ω_ref_, as can be seen via the manipulation:

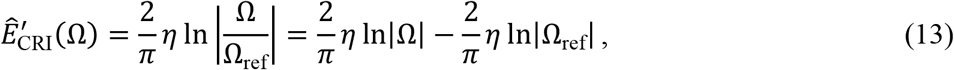 Thus, at Ω = Ω_ref_, the damper best approximates pure rate-independent damping, but in practice tuning Ω_ref_ can be complex. In §4.2 we discuss this tuning process in more detail.
ii. The time-domain formulation of this causal rate-independent damper can be derived through a memory-function approach, though we must pay careful attention to the singularity of the functions involved. Extending the analysis of Makris (1997a) using distributional methods (Estrada and Kanwal, 1989; Graf, 2010; Kanwal, 2004), we may express this memory function as:

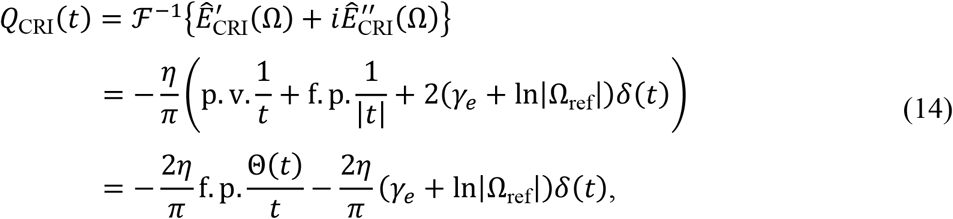

illustrated in Fig. 2. In Eq. 14 we introduce the Euler-Mascheroni constant *γ*_*e*_ ≈ 0.577, and two generalised functions: the Hadamard finite part regularisations (f. p.) of the singular ordinary functions 1/|*t*| and Θ(*t*)/*t*, where Θ(*t*) is the Heaviside step function, to distributions (Estrada and Kanwal, 1989; Kanwal, 2004; Monegato, 2009). In distributional literature, Θ(*t*)/*t* is often styled 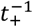. The f.p. regularisation in Eq. 14 means that, whenever we integrate or convolve these generalised functions, the integral is defined by Hadamard’s finite part: a process for assigning a finite value to diverging (*i*.*e*., infinite-valued) integrals by extracting the converging terms in a series expansion of the integral (Jones, 1996). This is necessary because integrals involving 1/|*t*| and Θ(*t*)/*t* often diverge—when so, they cannot be regularised via Cauchy principal value, but require stronger methods^†^. In the Supplementary Material (§S2), when we discuss practical numerical methods for evaluating finite part integrals, we provide a brief introduction to their theory.
iii. Convolving Eq. 14 with *x*(*t*) yields the complete time-domain formulation of the causal rate-independent damper, as the finite-part integral:

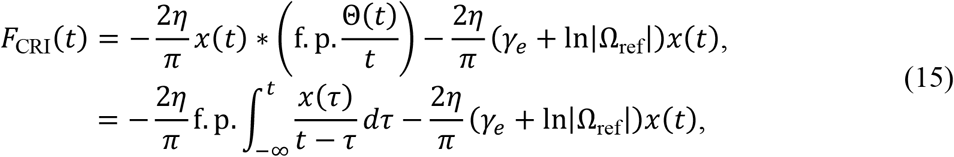 The effective stiffness—the coefficient of *x*(*t*)—may now be seen. This singular convolution integral is a causal variant of the Hilbert transform, with the upper limit of integration now *t*— ensuring causality, but at the cost of stronger singularity requiring finite-part evaluation.
iv. A practical problem with the causal rate-independent damper is that its stiffness becomes more and more unstable (negative-valued) as Ω drops below Ω_ref_. For any input or system dynamics known in advance, Ω_ref_ can be tuned to an appropriate frequency: however, the case Ω ≪ Ω_ref_ may arise if the system or input is not known in advance, or strong subharmonics are present in the signal. To guard against both these occurrences, we would be interested in limiting this negative stiffness, and this motivates a final approximate rate-independent model.

### 2.4. The Biot element

Causality analysis, as per Pons (2024), implies that *no* other linear causal models show perfectly rate-independent damping. The only pathway toward alleviating the negative stiffness of the linear causal model involves approximating rate-independence with some form of limited rate dependence. Several such approximations are available in structural dynamics and seismic analysis literature: from convenient but inaccurate approximations valid over only a narrow frequency band (Liu and Ikago, 2022; Reggio and De Angelis, 2015); to more complex generalised models which approach true rate independence (Eq. 12) in some parameter limit (Caughey, 1962; Luo and Ikago, 2021). Exoskeletal DMA data provides evidence for rate independence over a very large frequency range—up to three orders of magnitude (Dudek and Full, 2006; Gau et al., 2019)—and so we require a strong model. Here, we select the Biot element (Biot, 1958; Caughey, 1962; Luo and Ikago, 2021), formulated in the frequency domain as:

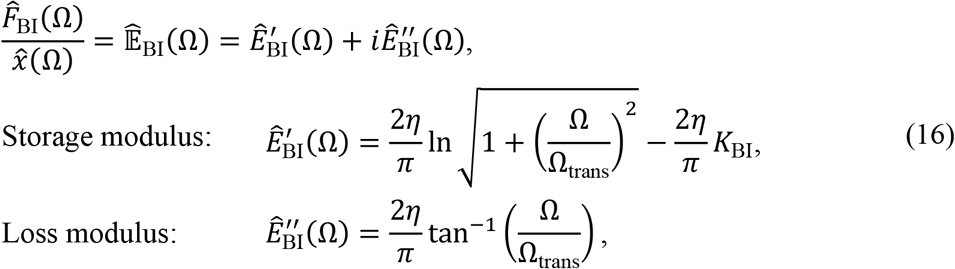

as per Fig. 2B. The parameter Ω_trans_ (> 0) is a transition frequency: the Biot element transitions between zero loss modulus at low frequency (|Ω| ≪ Ω_trans_) to near-rate-independent loss modulus at high frequency (at |Ω| ≫ Ω_trans_). The parameter *K*_BI_ is a stiffness corrector: if we wish to approximate rate-independent damping, Ω_trans_ is not a suitable operating frequency, because this is the frequency of transition. A more suitable choice is some higher Ω_ref_ = *N*_ref_Ω_trans_, where *N*_ref_ > 1 is a dimensionless factor describing how far above transition we wish to operate. However, at this Ω_ref_ the storage modulus is non-zero, and so we must select *K*_BI_ to correct this to zero. The correction value is:

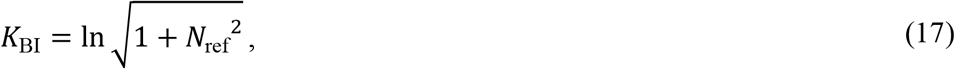

We suggest *N*_ref_ ≈ 20 for practical applications, as illustrated in Fig. 2B. Then at any given Ω_ref_, the loss modulus is 96% of its asymptote, but the storage modulus drops only to −4.2*η* at Ω = 0 (Fig. 2B). Note that selecting Ω_ref_ and *N*_ref_ defines Ω_trans_ for use in Eq. 16 and following. In this form, the Biot element approximates a causal rate-independent damper, and approaches it as *N*_ref_ → ∞ (*cf*. Fig. 2), but eliminates the pathology of infinite negative stiffness at Ω = 0.

Several novel practical properties of the Biot element may be derived. *First*, 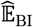 is equivalent to the principal branch of the complex logarithm (Ln) (Zill and Shanahan, 2009), providing the more concise formulation:

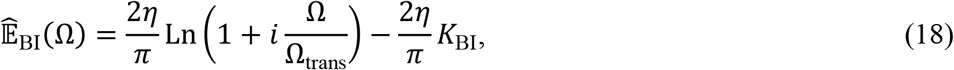

which appears to have previously gone unnoticed. *Second*, time-domain formulations of the Biot element have traditionally been expressed in terms of a convolution integral between the input velocity 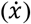 and the special function known as the exponential integral (Ei·) (Caughey, 1962; Luo and Ikago, 2021). However, there is again a simpler formulation—in terms of the element’s memory function, *Q*_BI_(*t*), which is given by:

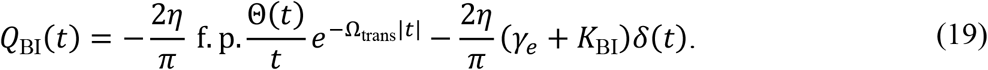

In the Supplemental Material (§S1) we present a brief derivation of this result, which to the author’s knowledge is presented for the first time here. Eq. 19 is visibly causal (Fig. 2A), and the general force response of the Biot element can thus be expressed:

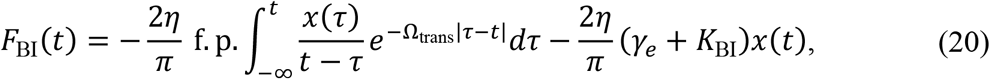

which again involves a strongly singular integral operator: a further variant of the Hilbert transform, with the causal integration limit of Eq. 15 and an additional exponential decay factor in the kernel.

Finally, we note that we have approached the Biot element as an approximation of the causal rate-independent damper, but it is also already known that the Biot element can be derived as an infinite sum of linear springs and viscous dampers, arranged in a generalized Maxwell configuration (Fig. 2C). Details are given in Caughey (1962) and Luo and Ikago (2021). Physically, this offers insight into one mechanism by which rate-independent damping can arise in an exoskeleton: as a bulk effect caused by a statistical distribution of elastic and viscous microstructures. We discuss this, and other mechanisms, in §6.1.

## 3. Computation and simulation

### 3.1. Computation of the Hilbert transform

For the time-domain formulations of §2 to be useful to insect biomechanics—whether in model identification or simulation—they must be convenient to compute for general numerical input data. Here we describe and develop numerical methods for computing these formulations. All methods presented across §3.1-3.2 are available as function code in MATLAB within the open dataset associated with this article—see the data availability statement.

To begin with a straightforward case: as a well-established singular integral operator, the Hilbert transform (Eq. 5), is the subject of many existing numerical implementations, in the form of the Discrete Hilbert Transform (DHT) (King, 2009; Marple, 1999). In MATLAB, the DHT is accessible via `hilbert()`; in Python, via `hilbert()` within SciPy; in R, via `hilbert()`within the gsignal package or `HilbertTransform()` within the hht package. Fig. 3 illustrates the performance of one such implementation (MATLAB DHT), for a triangle pulse and a sine wave—illustrative of limb perturbation (Dudek and Full, 2007), and flight motor oscillation (Pons and Beatus, 2022a), respectively. For both inputs, analytical solutions are available for validation. For the unit triangle pulse, as per King (2009):

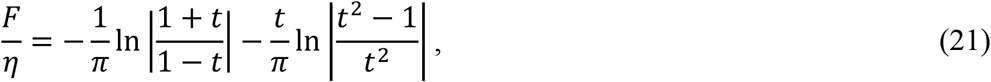

and for the sine wave, sin 2*πt, F*/*η* = cos 2*πt*, by the phase shift property. As can be seen in Fig. 3, MATLAB’s DHT routine is able to accurate capture these responses, and so we elide over the numerical details of this routine—excepting two practical notes:

**Figure 3.**
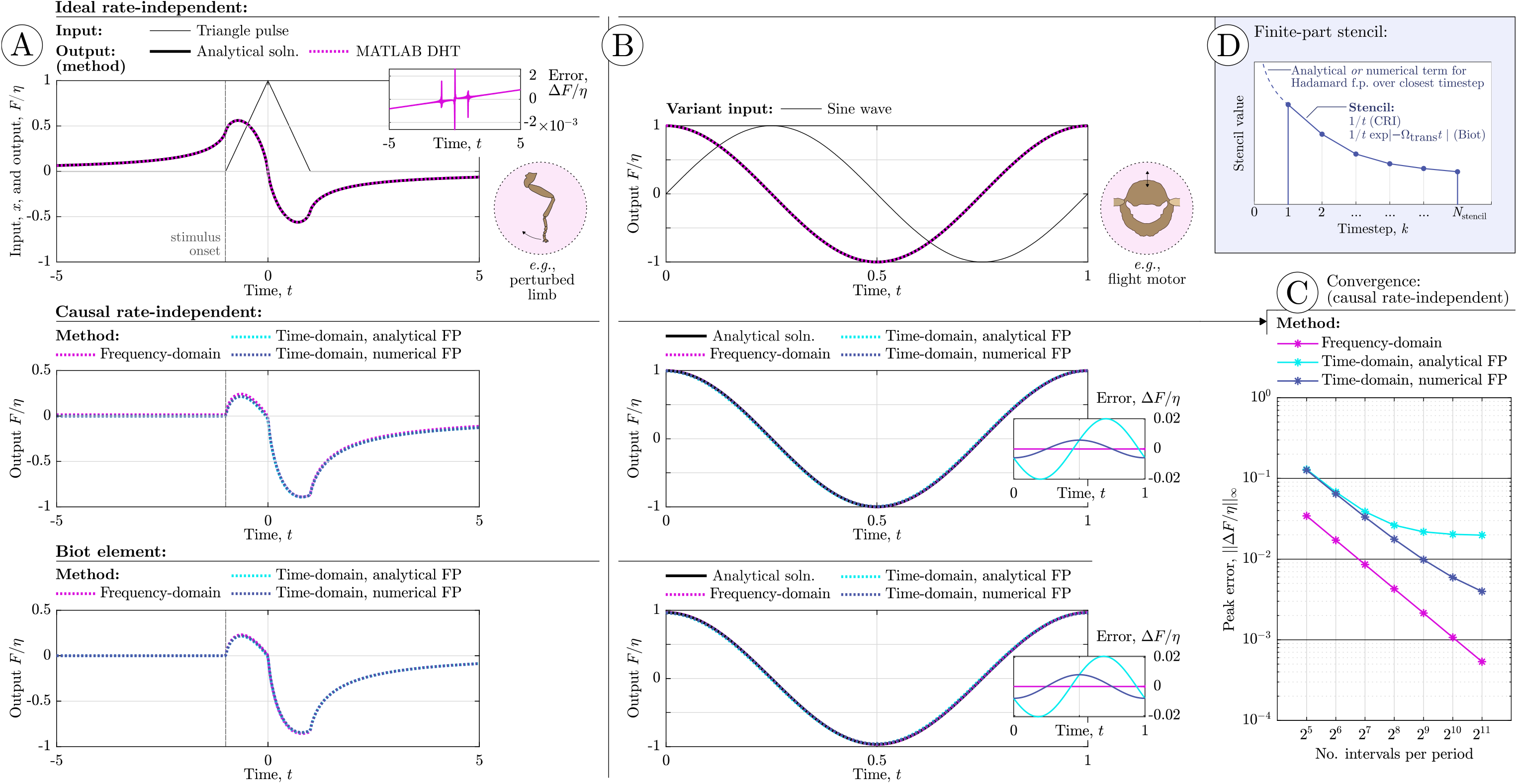
Performance, and illustrative results, of different numerical methods for computing the responses of the three rate-independent dampers (ideal, causal, Biot element). (**A**) Reponses to a triangle pulse input, representative of a limb perturbation, *cf*. Dudek and Full (2007) and Fig. 2. Analytical results are available for the ideal damper, via King (2009), permitting an error estimate. (**B**) Responses to a sine input, representative of flight motor operation, *cf*. Pons and Beatus (2022a). Analytical results are available for all three dampers, enabling an error estimate for the two causal dampers. (**C**) Convergence of time- and frequency-domain methods for the causal rate-independent damper: convergence is achieved, though slowly for the time-domain stencil, and this is limited by the (fixed) stencil length. (**D**) An illustration of the time-domain stencil for computing the finite-part integrals associated with the causal rate-independent damper and Biot element (Eq. 22): conventional quadrature stops at one timestep away from the present instant, and instead accounts for this closest timestep with a dedicated analytical or numerical approximation.

i. Because of the utility of the Hilbert transform in digital signal processing (DSP), many numerical routines, including all those above, output instead the *analytic signal*, of which the Hilbert transform is the imaginary part: *x*(*t*) + *i*ℋ{*x*(*t*)} (King, 2009; Marple, 1999). Hilbert transform results for structural modelling (Eq. 7) must be extracted as the imaginary part of this output.
ii. Because the Fast Fourier Transform (FFT) is computationally efficient, and the frequency-domain representation of the Hilbert transform is convenient and non-singular, many routines, including all those above, compute the Hilbert transform via the FFT (*e*.*g*., Marple, 1999). Time-domain quadrature methods are available (Gori and Santi, 1995), but not widely utilised. One effect of this choice, which is typically undocumented, is that these routines assume the input signal to represent a single period of a longer periodic signal (as per the FFT). This may be convenient in some contexts (*e*.*g*., periodic insect wingbeats), but inconvenient in others (*e*.*g*., perturbations of insect limbs). When non-periodic signals are under consideration, they must be zero padded, *cf*. Press (2007), before Hilbert transformation.

Finally, the comparison between ideal-damper responses to a triangle pulse and sine wave in Fig. 3 illustrate one wider practical point. The non-causality of the ideal damper is less observable for periodic inputs than for inputs involving an isolated stimulus. Qualitatively, the periodicity of the wave ensures that signals from the past and the future are similar and so the non-causality of the damper does not lead to significant changes in waveform, nor to a visible non-physical precursor response. Indeed, for a sinusoid (Fig. 3B), the waveforms for both causal and non-causal dampers are identical (another sinusoid). By contrast, when the input is an isolated event in time (*e*.*g*., a step or impulse), non-causality leads to an obvious non-physical anticipatory response (Fig. 2A). In biomechanical terms, this suggests that the ideal damper may be suitable for the analysis of steady wingbeat oscillation or steady walking; but not for isolated leg perturbations or strongly transient wing motion (*e*.*g*., takeoff, landing, and extreme manoeuvres).

### 3.2 Computation of finite-part singular operators

Computing the finite-part integrals governing the responses of the two causal dampers is more complex, and requires customised solutions. Broadly, two types of numerical methods are available: those performed in the frequency domain, via the FFT; and those performed directly in the time domain. Both have situational advantages and disadvantages. In this section we briefly present specific implementations of both types of method, and validate them in Fig. Implementations as function code in MATLAB are again available within the open dataset associated with this article.

#### Methods in the frequency domain

Frequency-domain computation of these causal damper responses is convenient and accessible. At some instant *t*, we: (**i**) compute the FFT of the input signal, 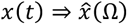; (**ii**) multiply by the damper’s complex modulus to obtain output force, 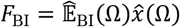 or 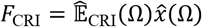; and (**iii**) inverse FFT back into the time domain, 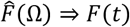. In the case of the causal rate-independent damper, this process contains a weak (logarithmic) singularity in the storage modulus, at 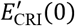. Because this weak singularity is integrable, it suffices to approximate 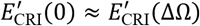, where ΔΩ is the frequency step of the FFT data. Because of the integrability of the singularity as ΔΩ → 0 (*i*.*e*., the input data *x*(*t*) gets finer) the damper’s response will converge, in the manner of a principal value (p.v.). As a corollary, it is also suitable to perform the computation with an even-number FFT spectrum, not containing Ω = 0, as this computation will also converge. In the case of the Biot element there is no singularity at all. Note that this mode of computation is distinct from frequency-domain analysis of linear systems—the computed results can be utilised in nonlinear analysis.

The weakness of the singularity in the frequency domain—compared to the strength of the singularity in the time-domain convolution—makes this mode of computation attractive, but it has the key disadvantage that the FFT always takes the input, *x*(*t*), to be periodic. If the damper is a component within a system that that is evolving according to its dynamics (*e*.*g*., an insect flight motor responding to muscle forcing), then extensive zero padding of the FFT computation is required to prevent bleed-over from the assumption of periodicity. To compute the impulse-like response in Fig. 3A, the triangle pulse is zero-padded for 20 pulse lengths on either side to mitigate bleed-over.

#### Methods in the time domain

Direct computation of casual damper responses in the time-domain is also possible via specialised numerical stencils—sets of weights over previous time points. Deriving these stencils requires deeper techniques for the analysis of finite-part integrals: these derivations are given in the Supplementary Material (§S2), alongside a brief introduction to the theory of finite-part integration. Here we present only the resulting numerical method. The responses of the causal rate-independent damper and Biot element may computed as:

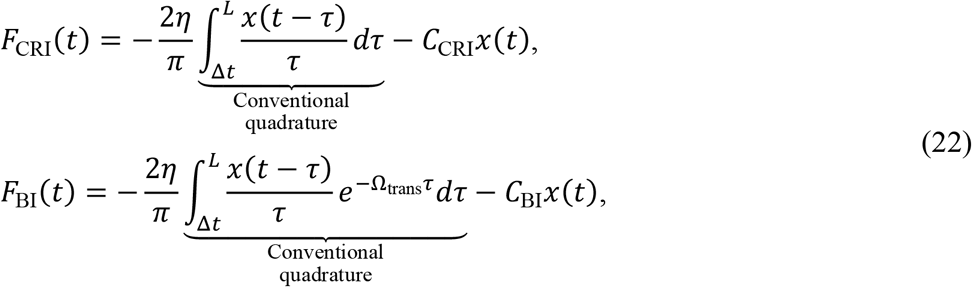

where the annotated integral term is now an ordinary integral which can be approximated by many conventional methods for numerical quadrature. In Fig. 3, and this article’s open dataset, Simpson’s 1/3 rule is used—the trapezoidal rule also works but converges slower; and adaptive Gaussian quadrature is known to be well-performing for these types of integrals, but is not directly consistent for sampled data with a fixed time interval.

Numerical parameters within Eq. 22 include the current timestep duration (Δ*t*), and the memory length of the damper (*L*), with larger *L* leading to more precise results. *C*_CRI_ and *C*_BI_ are corrector constants which depend generally on the current timestep duration (Δ*t*) and reference frequency parameters (Ω_ref_, *N*_ref_)—they represent the singularity that would otherwise be present in the integral, but has been extracted out (Supplementary Material, §S2), Two options for the computation of these corrector constants are available. *First* are analytical estimates, which are exact for the analytical model but not necessarily for the numerical implementation. These estimates are:

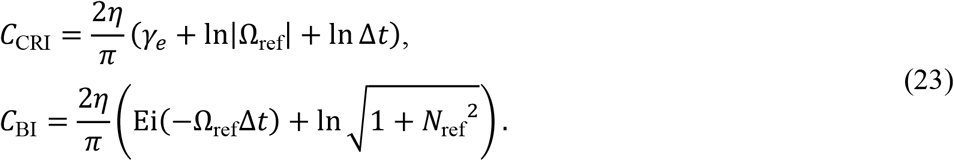

*Second*, numerical estimates for these constants are available, as the following single ordinary integral computations:

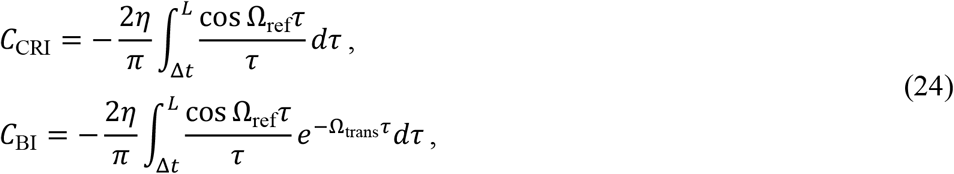

where, for the Biot element, Ω_trans_ is determined by Ω_ref_ and *N*_ref_. Practically: the purpose of defining Ω_ref_ is to ensure that the storage modulus of both dampers is zero at Ω_ref_. A direct way to ensure this is to compute the dampers’ responses to harmonic excitation at Ω_ref_ without any correction, and then identify the storage modulus within these responses and subtract this modulus as a corrector. Eq. 24 are simple computations of the uncorrected storage moduli at Ω_ref_, based on the fact that only the storage modulus, *not* the loss modulus, determines the dampers’ force responses at the displacement extrema of harmonic excitation (where, *e*.*g*., the velocity is zero). In Eq. 24, it is important that the same quadrature method is used as in Eq. 22, as these correctors can account for certain errors introduced via quadrature. In general, these numerical correctors are preferable: they are more accurate (Fig. 3C) and represent a trivial computational cost, equivalent to the computation of the response at a single timestep.

### 3.3. Forward simulation of exoskeletal structural models

A unique feature of the causal damping models is that they are compatible with structural models of the exoskeleton based on initial value problems in time. The causality of these models allows them to be simulated forward in time, with no knowledge of the future—as is typical for structural models, and as is representative of real exoskeletal behaviour. In this section we briefly outline simulation of a simple thoracic model with structural damping as an example. The model in question (Fig. 4A) is unforced and has linear mass (*M*), stiffness (*K*) and a Biot element with coefficient *η* = *Kγ*, where *γ* = 0.15 is roughly representative of lightly-damped exoskeletal structures (Gau et al., 2023; Wold et al., 2023). This model may be expressed:

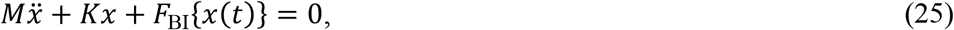

and has natural frequency 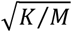, equivalent to natural period 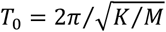. Discretising Eq. 25, with time vector **t** (elements *t*_*k*_, constant timestep Δ*t*), and displacement vector **x** (elements *x*_*k*_), we approximate the second derivative with a first-order backward difference in order to obtain a low-order explicit method for simulation. This yields:

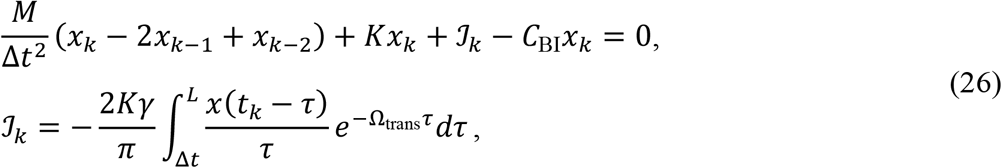

where ℐ_*k*_ denotes the computation of this ordinary integral by whatever quadrature scheme is chosen, and we use the analytical approximation for *C*_BI_ given in Eq. 23. Note that ℐ_*k*_ depends only on variables from the previous timesteps (*k* − 1, …), as *C*_BI_ accounts for any behaviour at *k*. Then the timestep can be advanced by computing *x*_*k*_ as:

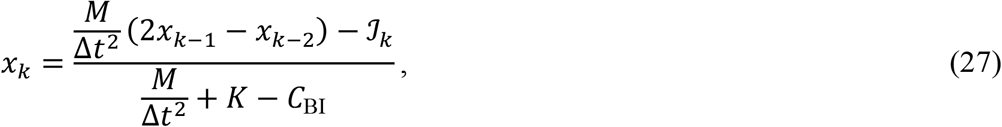

providing us with a simulation method for this causal system. Note that this numerical scheme is itself subject to numerical dissipation—for more precise simulations, variational approaches could be devised, *cf*. Pons and Cirak (2023). Fig. 4B-D illustrates the response of this model to an initial condition of *x* = 1 mm and 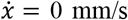, *e*.*g*., a long perturbation which is then suddenly released.

**Figure 4.**
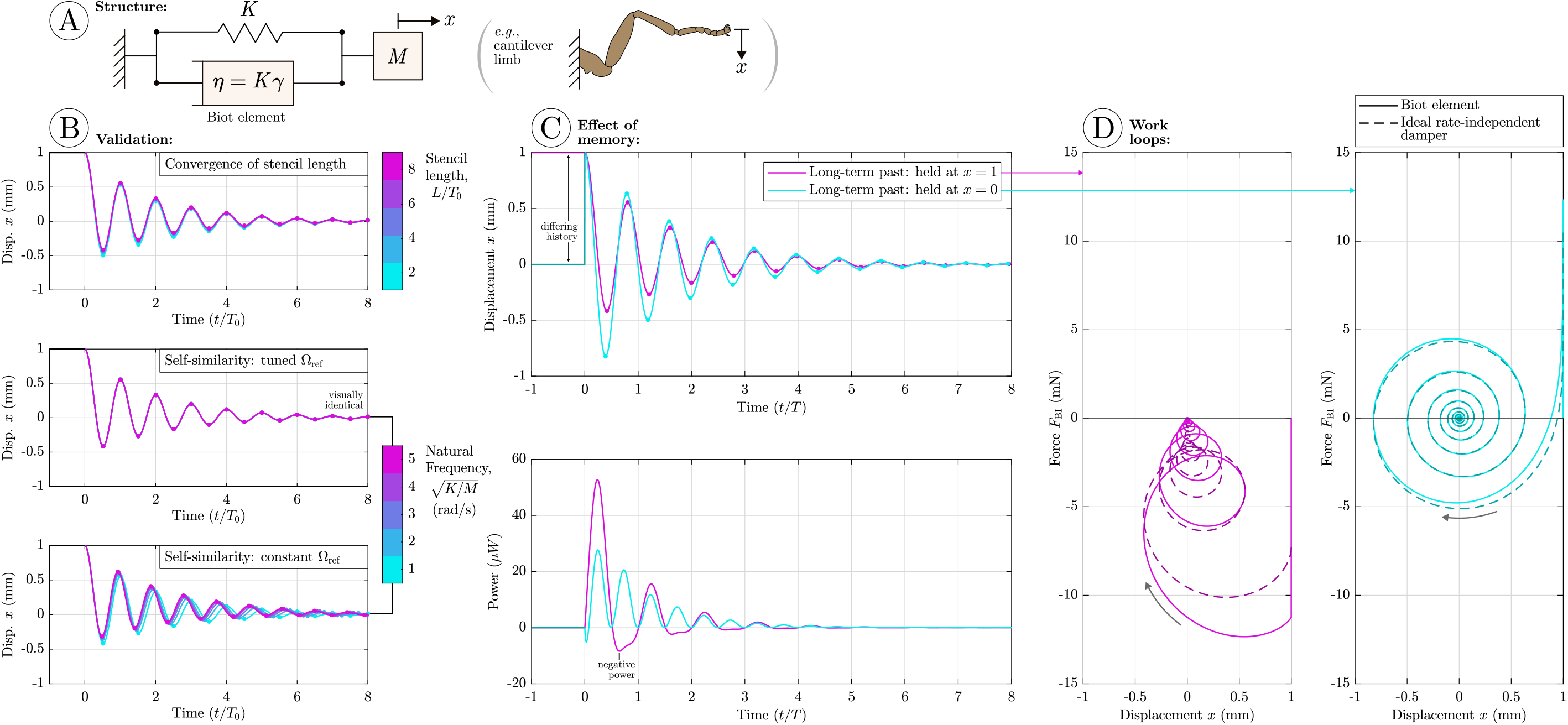
Forward simulation of the Biot element within an unforced spring-mass-damper system, representative, *e*.*g*., of a cantilever limb. (**A**) Free-body diagram: we consider *γ* = 0.15 and initial conditions *x* = 1 and 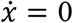. (**B**) Validation results: convergence under increasing stencil length (*L*) and self-similarity across varying natural frequency, indicating that the damped response does not vary with frequency (rate-independence). Exact self-similarity is observed when Ω_ref_ is tuned to each natural frequency; but only approximate self-similarity when it is held constant, due to the changing storage modulus (*cf*. Fig. 2B). (**C**) The strong memory of the Biot element has a significant effect: the long-term history of the system, before the initial condition, determines its response, and may lead to negative power generation, *i*.*e*., the damper doing work. (**D**) Work loops for the Biot element to both long-term histories, against the work loop for the ideal rate-independent damper (the Hilbert transform) under the same kinematics. The Biot element is a good approximation to the ideal rate-independent damper. However, both systems generate negative power in a way that indicates complex internal behaviour that may or may not respect the conservation of energy.

Two features illustrate that these simulation results are a good representation of causal rate-independent behaviour. *First*, the response of the damper is self-similar across frequency (Fig. 4B): when varying the model natural frequency, the response is scaled in time (by *T*_0_) but otherwise unchanged. This indicates that the damping within the model is frequency-independent. In the case where Ω_ref_ is tuned to each natural frequency the self-similarity is exact; in the case where it is held constant the self-similarity is approximate, due to the changing storage modulus as the response frequency departs from Ω_ref_ (Fig. 2B). *Second*, the damper forcing, ℐ_*k*_ − *C*_BI_*x*_*k*_, is a good approximation to the forcing of the ideal rate-independent damper (Fig. 4D). This indicates that the scale and waveform of forcing is representative of ideal rate-independence.

In performing these simulations, we reveal several interesting and concerning properties of the Biot element. Because the force response of the Biot element is an integral over past time, the entire system’s response is strongly dependent on its long-term history, *i*.*e*., its behaviour before the initial condition. Fig. 4C compares this long-term memory effect for two prescribed long-term histories: the system held at *x* = 1 mm over an infinite past, and the damper held at *x* = 0 mm and then suddenly brought to *x* = 1 mm right before the system is released. These two histories lead to free oscillations with large differences (∼100%) in oscillation amplitude. As we discuss in §6.2, this presence or absence of memory effects in real exoskeleta is one characteristic which may help filter between differing models of rate-independent damping.

However, more concerningly, in some circumstances, this memory effect can lead to responses with strange energetic properties. Fig. 4C also compares the instantaneous power dissipation of the Biot element 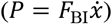 under the two system responses. Both responses contain regions of negative power (*P* < 0), representing instantaneous power *output* from the damper. Here, there is a challenging paradox to resolve. In a passive structure, negative power does not necessarily violate energy conservation, provided that this energy is stored and released via a conservative potential, *i*.*e*., structural elasticity. From the work loops in Fig. 4D, we see that this negative power is *not* explained by any single linear or nonlinear parallel or series elasticity, as per Pons and Beatus (2022b, 2023): considered as an SDOF system, the Biot element appears to violate the conservation of energy. However, this element can also be constructed from an array of dissipative and elastic elements (Fig. 2C) which should not be capable of this violation. The apparent resolution to this paradox is that, energetically, the Biot element behaves an MDOF rather than SDOF structure, with internal elements of energy storage that depended not only on instantaneous *x* but on its complete history. This leads us to the topic of §4.

## 4. Energetic properties of rate-independent damping

### 4.1. Non-dissipative behaviour of the ideal rate-independent damper

In §3.3, we observed that, as we computed the energy dissipation of the Biot element under the free oscillation of an exoskeletal model, this element started to output energy over certain time windows, rather than dissipate it. This is a concerning feature, as these models are, by definition, intended to model a source of energy dissipation within the insect exoskeleton. To understand this issue further, we return to the ideal rate-independent damper (§2.2), which has not storage modulus and thus should not store energy. Under pure simple harmonic motion, this damper dissipates energy at all times: input *x* ∝ sin *t* yields output *F* ∝ cos *t* with power dissipation 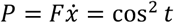 which is ≥ 0 always. But, to the author’s knowledge, it has not previously been observed that these dissipative properties do hold not for general motion— just as all conceivable loss moduli are not necessarily causal (Pons, 2024), all conceivable loss moduli are not necessarily always dissipative. This can be demonstrated by a simple counterexample. Consider motion with just two harmonics:

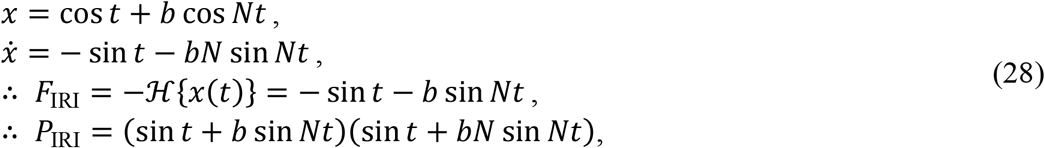

*P*_IRI_ is illustrated for *N* = 6 and *b* = 0.5 in Fig. 5A. Regions of negative power are clearly observed. Whether this negative power violates the conservation of energy is a challenging question that we cannot completely resolve. This ideal damper has no storage modulus: in physical terms, it is incapable of storing energy under any kind of sinusoid excitation. In addition, as this damper is non-causal, it cannot be represented in the MDOF generalized Maxwell configuration of Fig. 2C. Both these points indicate that this negative power likely violates conservation of energy—but we cannot rule out the possibility that some kind of non-causal conservative MDOF assembly exists^‡^.

**Figure 5.**
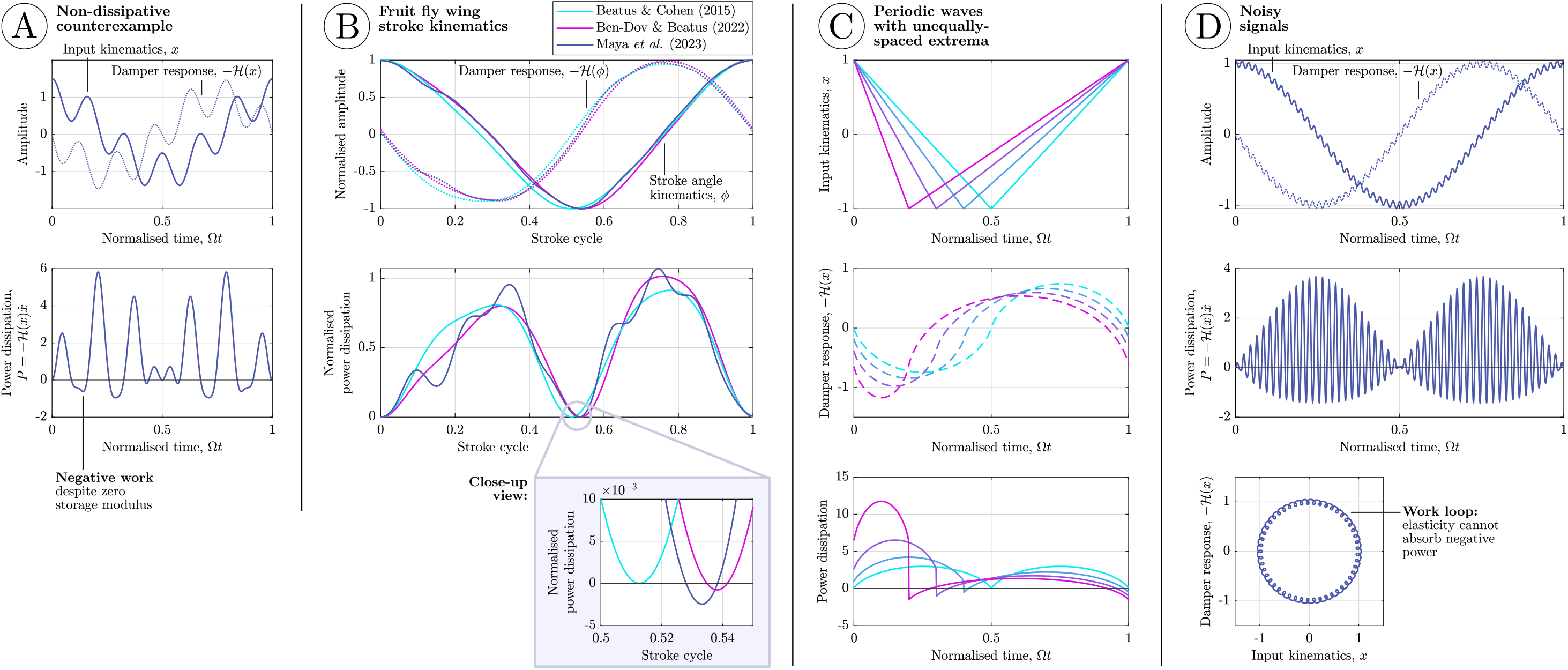
Energetic properties of the ideal rate-independent damper—a short catalogue of responses to additional waveforms. (**A**) The initial non-dissipative counterexample, Eq. 28, in which negative power is observed. (**B**) Wingbeat kinematics from the fruit fly *Drosophila melanogaster*: stroke angle under hovering flight, from several sources (Beatus and Cohen, 2015; Ben-Dov and Beatus, 2022; Maya et al., 2023). Practically, these wingbeat kinematics are completely dissipative; but strictly, a small level of negative power is present (≈0.1% of peak power). (**C**) In periodic waves, unequal spacing of the displacement extrema in time can lead to non-dissipative behaviour—the case of a triangle wave is illustrated, partly explaining the negative power in (B). The level of negative power introduced by this effect is small. (**D**) The case of noisy signals, illustrated with a deterministic two-harmonic signal, analogous to (A). The ideal rate-independent damper generates negative power when exposed to noise. Observing the work loop, we can confirm that this negative power cannot be explained by a SDOF nonlinear elasticity, *cf*. Pons and Beatus (2022b; 2023): either internal DOFs are present, or conservation of energy is violated.

Practically, however, this counterexample indicates that the ideal rate-independent damper can be non-dissipative: under certain inputs, it generates negative power. Previous frequency-domain implementations (as *iγ*) have assumed this structure to be a damper (Dudek and Full, 2006; Pons and Beatus, 2022a; Wold et al., 2023), but this assumption is unsafe. *In lieu* of a deeper theoretical characterisation, we provide a short catalogue of the damper’s behaviour across several key waveforms found in insect biomechanics—as follows:

i. The triangle pulse waveform in Fig. 2-3 is purely dissipative. This applies to other isolated symmetric pulses, *e*.*g*., Gaussian, cosine, and square pulses; but not to asymmetric versions of these pulses (half-Gaussian, half-cosine, *etc*.). Conveniently, these properties can be observed directly in tabulated Hilbert transform results (King, 2009): for pure dissipation, the Hilbert transform and pulse velocity show opposite sign always.
ii. Dipteran wing stroke angle kinematics, being close to sinusoid, are dissipative for practical purposes. Figure 5B illustrates this for several recent estimates of the wing stroke kinematics for hovering *Drosophila melanogaster* (Beatus and Cohen, 2015; Ben-Dov and Beatus, 2022; Maya et al., 2023). Strictly, however, these kinematics show a very small level of negative power—approximately 0.1% of peak positive power.
iii. When a periodic wave has displacement extrema are unequally spaced in time, non-dissipative behaviour can arise. Figure 5C illustrates this effect for a triangle wave. In an insect flight motor, this unequal spacing represents, *e*.*g*., the upstroke being slightly faster than the downstroke, as in *D. melanogaster* —partly explaining the negative power in (**ii**) and Fig. 5B. However, the negative power introduced by this effect is small, and the rate-independent damper may remain a reasonable approximation.
iv. Noisy motion is *not* dissipative (Fig. 5D)—the addition of noise to any signal leads to non-dissipative noise in the response, as this noise behaves essentially as high-frequency multiharmonic components, *cf*. Fig. 5A. The effect can be removed by filtering noise out of the signal, but notably this means that rate-independent dampers cannot effectively model a structure’s response to noise excitation—*e*.*g*., the white noise motion that Wold et al. (2023) applied to hawkmoth thoraces.

### 4.2. Energetic tuning of the reference frequency in causal dampers

The situation in the causal rate-independent damper and Biot element is more complex, as these structures have a nonzero storage modulus which itself will contribute to instantaneous energy generation in a way which may or may not be consistent with conservation of energy (true energy storage *vs*. energy generation). In these structures we also have a reference frequency, Ω_ref_, as a free parameter. The role of Ω_ref_ can be seen as an energetic one: its purpose is to tune the damper to have zero storage modulus, *i*.*e*., zero negative power, at a certain frequency. Broadly, the classes of energetic behaviour identified in §4.1 apply to these dampers also, as this behaviour arises as a result of the constant loss modulus. However, there is one further energetic effect that is worth isolating. For sinusoidal motion at frequency Ω, zero storage modulus, and thus no negative power, is achieved by selecting Ω_ref_ = Ω: a transparent energetic tuning relationship. However, for periodic waves at fundamental frequency Ω that are *not* purely sinusoidal, the value of Ω_ref_ that exactly ensures no negative power (if such a value exists) is generally not exactly equal to Ω.

Figure 6 illustrates this phenomenon for the response of a causal rate-independent damper (§2.3) to a symmetric triangle wave. Denoting Ω_0_ the fundamental frequency of this triangle wave, a state of pure dissipation is reached at Ω_ref_ ≈ 1.39Ω_0_. If we were to select Ω_ref_ = Ω_0_ regardless, negative power would be present, to the level of ≈26% of the peak positive power and ≈0.35% of the overall absolute work. While this would be a noticeable peak for this strongly non-sinusoidal triangle wave, most insect wingbeat kinematics are not so extreme (*e*.*g*., Fig. 5B) and so Ω_ref_ = Ω_0_ may be a reasonable approximation. However, the fundamental principle still holds: that the appropriate Ω_ref_ depends not only on the frequency of a wave but *also* its waveform. One practical implication of this is that tuning Ω_ref_ directly to the under identification will lead to more accurate results than assuming this frequency based on the dominant frequency within the data, particularly with regard to the damper’s energetics.

**Figure 6.**
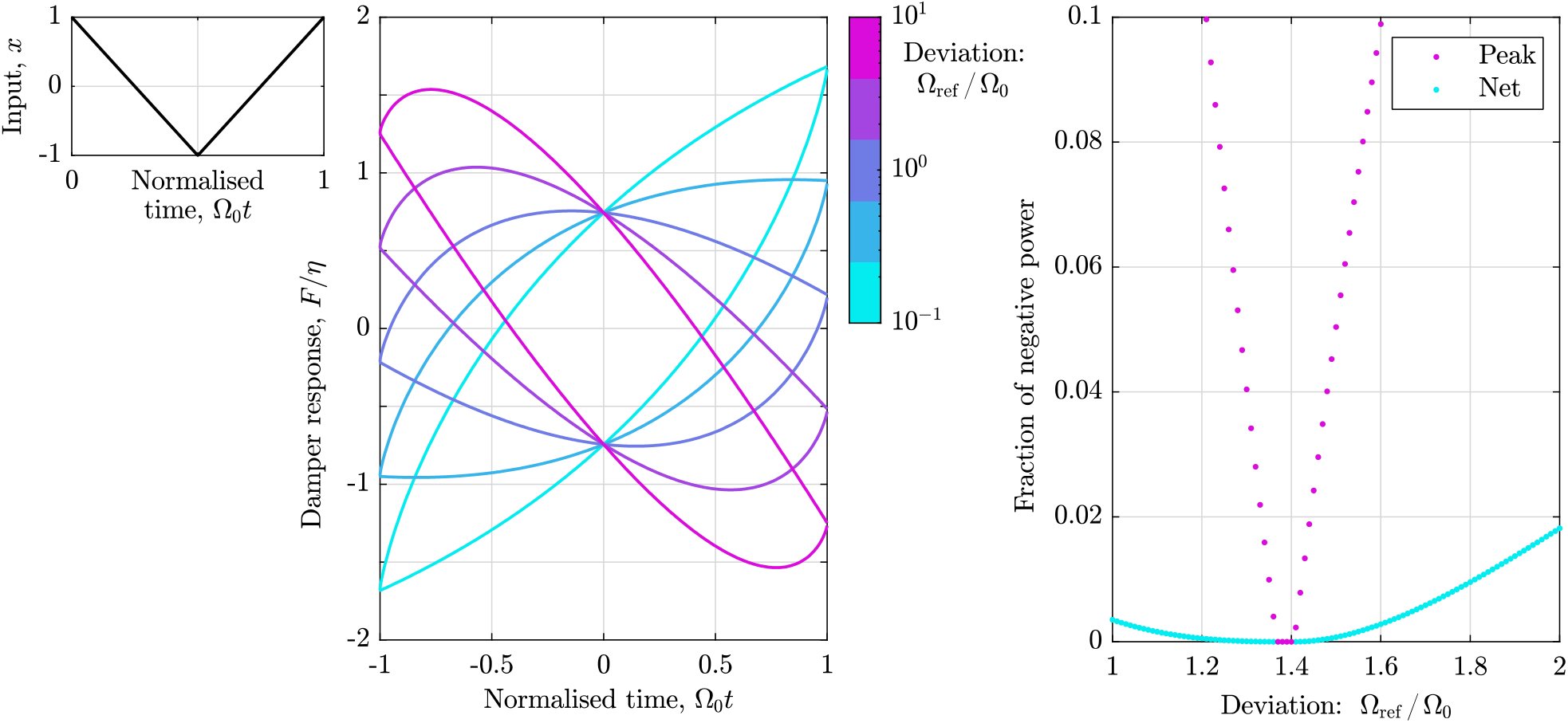
Energetic tuning of the causal rate-independent damper. The effect of selecting Ω_ref_ ≠ Ω_0_ for a triangle wave at fundamental frequency Ω_0_ is shown. The work loops of the damper’s response under Ω_ref_ illustrates that the negative power generated by Ω_ref_ can be completely accounted for by elasticity: a midline elasticity exists, as per Pons and Beatus, (2022b; 2023). However, Ω_ref_ = Ω_0_ does not lead to no negative power: instead, this state is reached at Ω_ref_ ≈ 1.39Ω_0_. Away from this point, the level of negative power—measured as peak fraction, and net (absolute integral) fraction—increases.

## 5. Application to exoskeletal modelling

### 5.1. Time-domain identification based on exoskeletal DMA data

Finally, with the results of §2-4, we can make progress on several challenging problems in insect biomechanics that pertain to rate-independent damping. The first challenge is raised by Gau et al. (2019) and Wold et al. (2023) in the context of rate-independent damping within the exoskeleton of hawkmoths (*Manduca sexta*) within flight. The challenge is that existing frequency-domain methods for identifying rate-independent are restricted to a very specific form of data: loss moduli, measurable when a structure that behaves linearly is subjected to sinusoidal excitation during DMA. If loss moduli cannot be estimated due to the nonlinear effects—*e*.*g*., nonlinear elasticities, as per Gau et al. (2019)—then frequency-domain identification breaks down. And if we wish to study the case of non-sinusoidal excitation, as per Wold et al. (2023), to better understand thoracic damping effects during insect flight manoeuvres, then frequency-domain identification again breaks down—and worse, we have no method of confirming whether a rate-independent damping model remains valid.

These challenges are solved by the time-domain formulations in §2. In Fig. 7, we demonstrate fitting a rate-independent damper with a nonlinear elastic term to DMA data for a hawkmoth exoskeleton under non-sinusoidal ‘asymmetric’ excitation (Wold et al., 2023). The fitted model is:

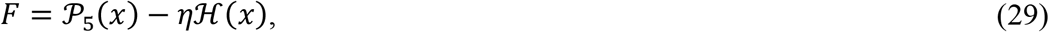

for force *F* (mN) and displacement *x* (mm). This time-domain model represents an ideal rate-independent damper, −*η*ℋ(*x*), in parallel with a nonlinear elasticity defined by a fifth-order polynomial in *x*, 𝒫_5_(*x*), *i*.*e*.:

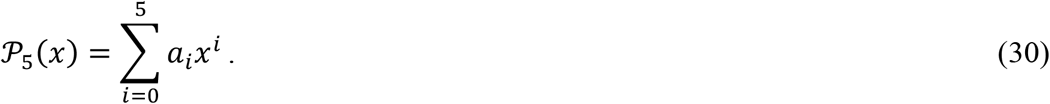

**Figure 7.**
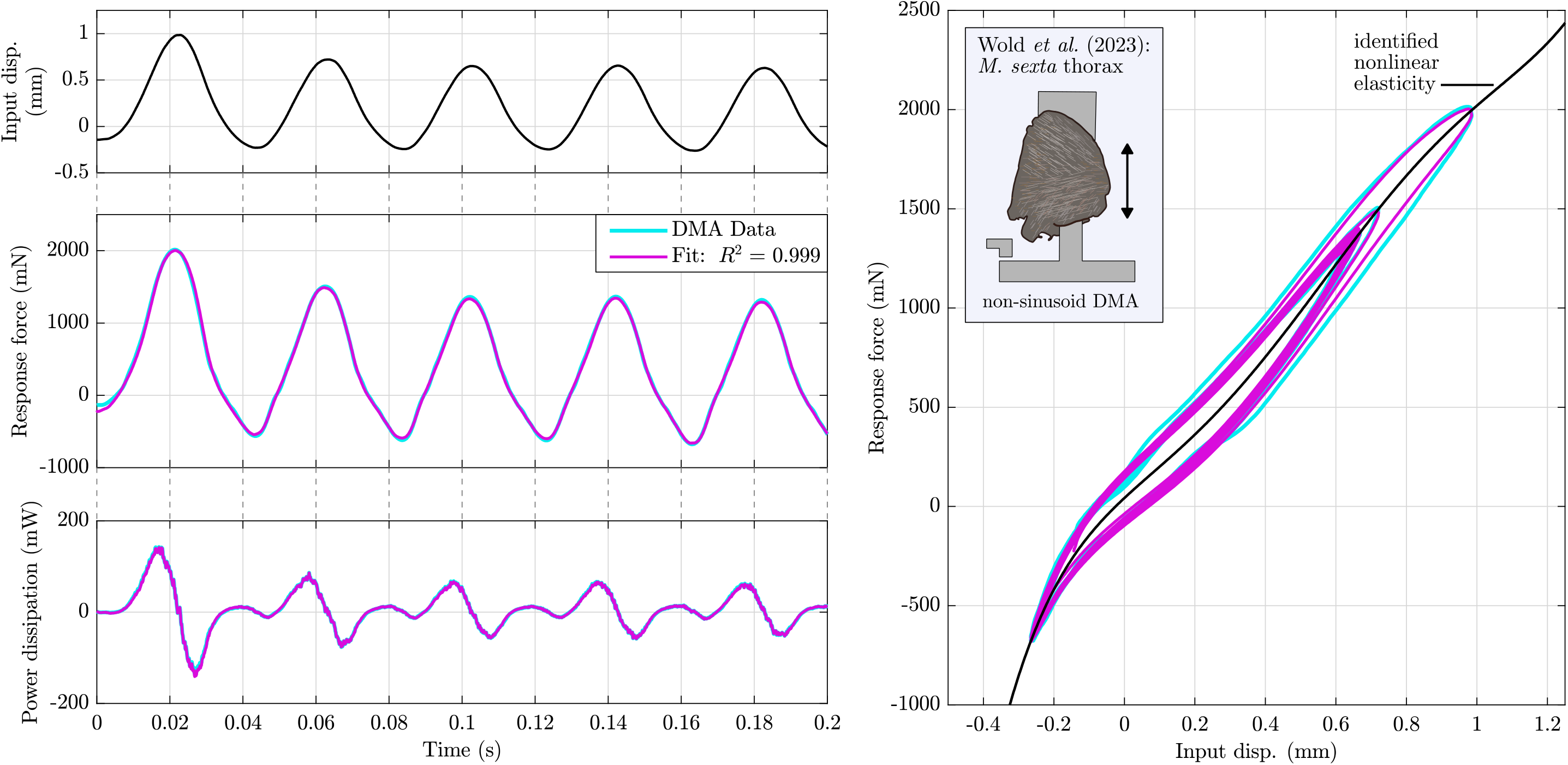
Time-domain identification of rate-independent damping in the asymmetric DMA data of Wold et al. (2023), for a *Manduca sexta* thorax, accounting for nonlinear elasticity up to fifth-order polynomial (Eq. 29). Illustrated is a sample interval, 0 ≤ *t* ≤ 0.2 of the DMA input displacement signal, the response force data and fitted profile, and the response and fit power dissipation; as well a response and fit work loop over the complete fit interval, 0 ≤ *t* ≤ 1. Note that the displacement and force data are subjected to Savitsky-Golay smoothing (5^th^ order over a frame width of 5.1 ms) in order to eliminate high-frequency non-dissipative noise (§4.1); and that the remaining low-frequency negative power arises from energy storage within the nonlinear elasticity, 𝒫_5_(*x*).

Fit parameters are *η* and the elastic parameters *a*_0_-*a*_5_. Fitting is performed in the time-domain using a least-squares method; Fig. 7 illustrates the fitted result in time and in a work loop. We obtain an accurate fit (*R*^2^ = 0.999), indicating that a rate-independent damping is suitable, at least for this individual time series. We estimate *η* = 277 N/m, and if we define a dimensionless damping term *γ* as *η* = *a*_1_*γ*, then this corresponds to *γ* = 0.167, appropriate for the hawkmoth thorax (Wold et al., 2023). This is an illustrative solution to the challenge posed by Wold et al. (2023): it not only allows us to accurately study the case of non-sinusoidal input signal, but also to account for the significant nonlinear elasticity in the exoskeleton—nonlinearity observable in the data and fit work loop (Fig. 7). Further application to the full dataset of Wold et al. (2023) would lead to further insight into the suitability of rate-independent damping as a model of structural dissipation within the insect flight motor.

### 5.2. Time-domain identification based on transient exoskeletal responses

A second challenge, extending that covered in §5.1, is posed by Dudek and Full (2007): the identification of rate-independent damping based on the free response of an insect limb—here, a metathoracic limb of the deathhead cockroach *Blaberus discoidalis*—following a perturbation. This involves identification of a complete oscillator model, involving rate-independent damping, elasticity, and mass, from the perturbation response. The approach of Dudek and Full (2007) to this identification is to assume an oscillator configuration, involving am ideal rate-independent damper, perform a suite of approximate forward oscillator simulations, ignoring the noncausal signal, and then identifying model parameters by matching the forward simulation results. However, there is an alternative inverse approach that is more efficient and more generalisable. Extracting the free displacement response data, *x*(*t*), we can compute a library of oscillator component force terms: elasticity, *x*(*t*); ideal rate-independent damping, −ℋ{*x*(*t*)}; and inertia, 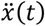. For a linear oscillator:

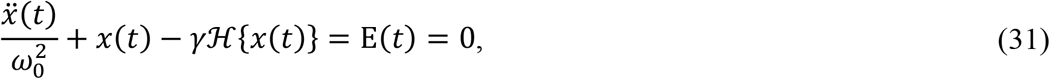

where we have two unknown parameters—natural frequency ω_0_, and dimensionless damping *γ*—and a residual E(*t*) that should be zero for an exact fit. We subject *E*(*t*) to least-squares minimisation and identify ω_0_ and *γ* directly: an identification based on forward simulation of the oscillator model, but on its internal consistency.

Figure 8 illustrates this process for a sample free response profile, that reported by Dudek and Full (2007) in their Fig. 2B. As an illustration of goodness-of fit, we may observe the fitted residual E_fit_(*t*), which has the same units as displacement; and may also define a fitted displacement profile as:

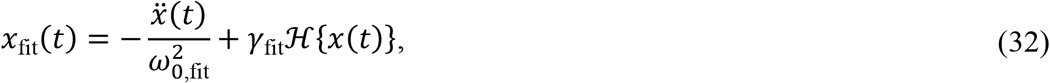

illustrated in Fig. 8. We identify *γ* = 0.52, matching but slightly lower that the estimate of Dudek and Full (2007) for the same profile (*γ* = 0.58). This approach is an illustrative solution to the challenge of full oscillator identification from kinematic data: it is (**i**) is faster and more accurate than attempting to simulate a causal (or non-causal) oscillator in the time domain, as it requires no forward simulation and thereby completely avoids numerical dissipation; and (**ii**) can easily account for arbitrary nonlinearities that are hard to simulate, *e*.*g*., a polynomial elasticity, which can be added directly to Eq. 31 with no change in identification process. Further application to a wider dataset would again lead to further insight into the suitability of rate-independent damping as a model of insect limb energy dissipation.

**Figure 8.**
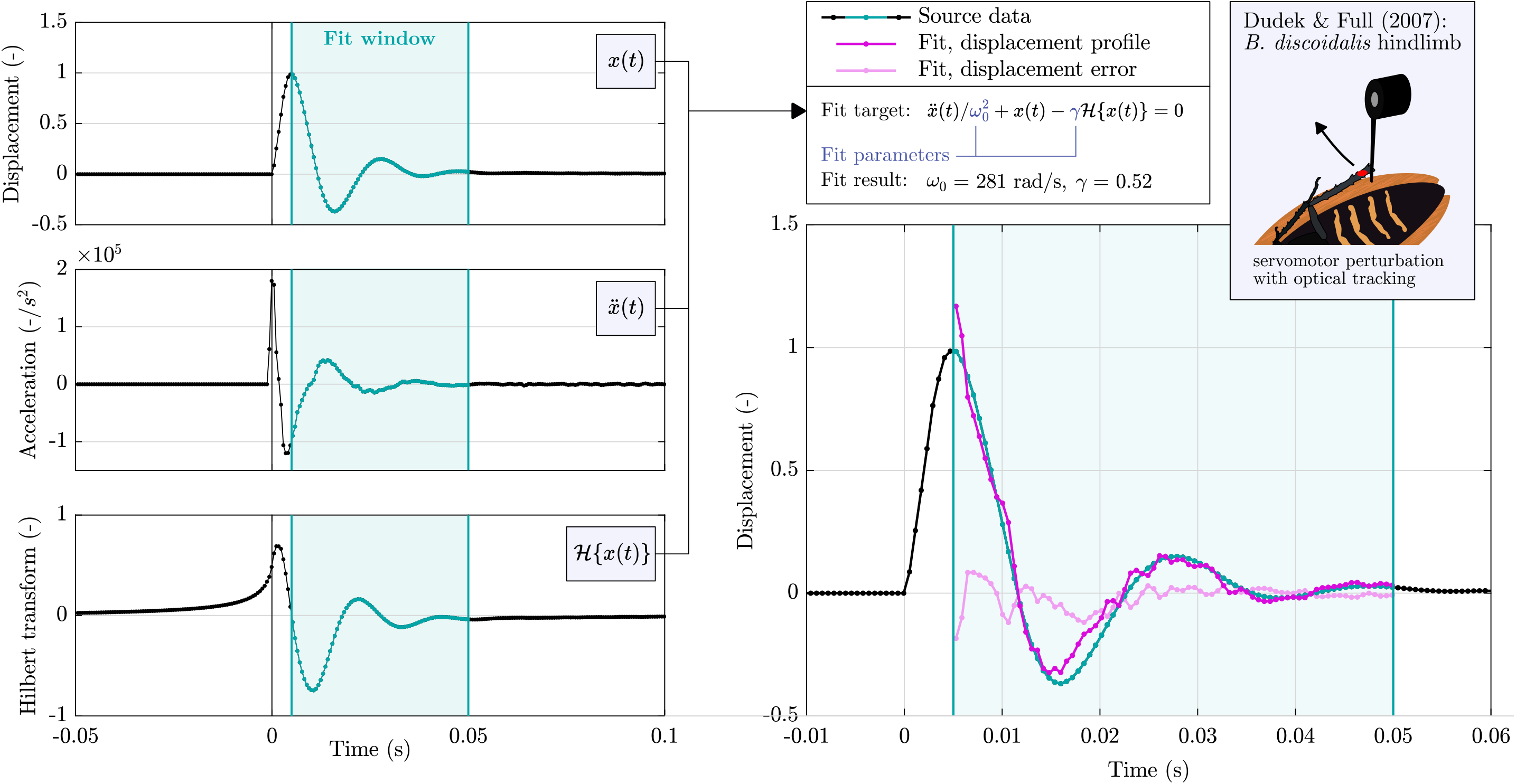
Time-domain identification of a linear oscillator with ideal rate-independent damping in the perturbation data of Dudek and Full (2007), for a hindlimb of *Blaberus discoidalis*, via an inverse approach. Illustrated are the component time signals, *x*(*t*) and the derived 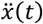 and ℋ{*x*(*t*)}; and the fit results, E_fit_(*t*) and *x*_fit_(*t*). Note that, as per the data in Dudek and Full (2007), displacement is in normalised units. We select a sample window starting after the displacement extrema: this results in a good fit, matching Dudek and Full (2007), whereas fitting prior to this point is poor, either because the servomotor is still in contact with the limb; the acceleration signal data is too coarse; and/or the damper and linear stiffness model breaks down.

### 5.3. Simulation of full flight motor systems

The final challenge which is partly solved by these time-domain methods is that raised by Lynch et al. (2022) and then Gau et al. (2023): the challenge of forward simulation of insect flight motors, under asynchronous muscle forcing, accounting for exoskeletal rate-independent damping. Forward simulations of these integrative models of insect flight motors have helped reveal deep properties of these motors, including their evolutionary history (Gau et al., 2023); but structural damping, *e*.*g*., as identified in §5.1, has been a missing component. In §3.2, we demonstrated methods for forward simulation of full oscillator models with causal rate-independent damping, and so solved this challenge at a methodological level—the numerical method of Eq. 26-27 is applicable to a linear flight motor model, and easily extensible to the nonlinear case. However, while these time-domain formulations give us the ability to simulate causal rate-independent damping, they also highlight that care is needed when doing so. Outside of a narrow set of input signals, rate-independent damping shows concerning energetic behaviour: negative power generation (§4.1) as well as counterintuitive tuning relationships (§4.2). These phenomena pose a particular risk in forward simulation, as the damper input signal is not known *a priori*, and negative power generation by the damper may have a long-term impact on the system response. At the least, robust post-processing checks are required to ensure physical behaviour—but we suggest, further, that that frequency- or rate-independent damping as currently formulated in biomechanical literature is fundamentally unsuited to general forward simulation. However, our analysis thus far offers insights into the development of more general accurate general models—we consider this in §6.

## 6. Discussion and conclusion

### 6.1. Origins of rate-independent damping

That rate-independent damping of some form is present is the insect exoskeleton is borne out by experimental data (Dudek and Full, 2006; Gau et al., 2019; Wold et al., 2023); but the physical mechanisms underlying it remain unclear. Several non-exclusive hypotheses can be made, and these hypotheses align in a revealing way with different models of rate-independent damping we presented in §2.

The *first* hypothesis is that this damping is a manifestation of structural hysteresis arising from elastoplastic behaviour: a transition between recoverable (elastic) and irrecoverable (plastic) deformation as strain increases (Giorgi and Morro, 2021; Zheng, 2019). In metals, the grain-related mesoscopic mechanisms behind this transition are well-studied (Weng, 1983; Zheng, 2019). Harder materials within the exoskeleton may show elastoplastic behaviour: there is evidence for plasticity both the chitin-polymer matrix of insect cuticle (Hillerton et al., 1982) and in chitin itself (Mir et al., 2008), but details are unclear. Elastoplasticity generates true quasistatic hysteresis: energy dissipation is maintained in the quasistatic limit, *i*.*e*., as Ω → 0. In a linear model, this corresponds to a constant loss modulus as Ω → 0, which, following §§2-5, we know to be physically impossible: such a linear model is necessarily non-causal and/or shows unstable negative stiffness and/or violates conservation of energy. This implication here is that true hysteresis is beyond the reach of linear analysis: it is a nonlinear phenomenon which is impossible to linearise without sacrificing basic physical principles.

The *second* hypothesis is that this damping represents a viscoelastic effect that is largely rate-independent but remains fundamentally a *dynamic* rather than hysteretic effect. Here, the Biot element (§2.4) offers the insight that rate-independent damping can arise from an ensemble of mesoscale viscous dampers and linear elasticities (Fig. 2C). This process is analogous to how with weakly rate-dependent (fractional-order) damping can arise from a hierarchy of self-similar mesoscale elements within vertebrate cartilaginous tissues (Guo et al., 2021). In the exoskeleton, there are several possibilities for this mesoscale ensemble, extending from the mesoscopic composition of cuticle to the varying thickness and composition of the sclerites (Barbakadze et al., 2006; Casey et al., 2022). Yet, at core, this viscoelastic damping is still a fundamentally dynamic process: it cannot show dissipation under true quasistatic motion, and the loss modulus must tend to zero as Ω → 0. In this sense it is a fundamentally different process to true hysteresis, and *can* be modelled linearly, as per the construction and properties of the Biot element.

Distinguishing between these hypotheses does not appear possible with current data, and indeed, both effects may be present in the insect exoskeleton. Model-identification approaches may be an effective approach to resolve between them: each effect has naturally associated models (nonlinear hysteretic *vs*. linear viscoelastic), and testable predictions are available for each. We discuss these topics in §6.2.

### 6.2. Alternative models and testable predictions

We began this study with the classical frequency-domain exoskeletal damping model: the complex stiffness *iγ*. As time-domain formulations reveal, this model is only realistic under harmonic excitation: outside this narrow condition, it observably violates causality and can generate negative power. Studying rate-independent damping under general exoskeletal motion requires alternative models—and in this work, we have already suggested two: the causal rate-independent damper and the Biot element. Future prospects for general models of rate-independent damping are inflected by the two hypotheses for its origin that we outlined in §6.1.

In the direction of models for general *viscoelastic* rate-independent damping, the Biot element is a promising candidate. Fractional-order models (Luo and Ikago, 2021) are an alternative with more pronounced rate scaling (*i*.*e*., weaker rate-independence), and are currently used in structural models of insect asynchronous muscle (Swank et al., 2006). Considered as a class, these viscoelastic models have a common characteristic: they show a storage modulus that is varies with frequency. This varying storage modulus is a key testable prediction of viscoelastic modelling, and an understudied feature in existing structural damping studies. If linear viscoelastic damping is present, then a varying storage modulus *will* be observed: and if this damping is approximately rate-independent, then the storage modulus scaling will be approximately logarithmic (§§2.2-2.3). Indeed, evidence for varying storage modulus may already be available: the raw time-series force data of Gau et al. (2019) shows increasing peak force with frequency, which may indicate an increasing thoracic storage modulus.

In the direction models for general *hysteretic* rate-independent damping, the overwhelming consensus of our analysis is that linear hysteresis is non-physical: nonlinear modelling is required. Studies of structural hysteresis in other materials, such as rock (Guyer et al., 1995) and rubber (Penas et al., 2022), suggest several directions for nonlinear modelling, including thermodynamics-driven constitutive modelling with relaxation functions, and assemblies of hysteretic mesoscopic elements, or *hysterons* (Bertotti and Mayergoyz, 2006; Giorgi and Morro, 2021; Guyer et al., 1995). A key testable prediction of these models, and of quasistatic hysteresis generally, is the phenomenon of endpoint memory (Claytor et al., 2009; Vakhnenko et al., 2005), otherwise known as return-point memory (Zirka and Moroz, 1999), or Madelung’s second rule (Brokate and Sprekels, 1996; Penas et al., 2022): the material remembers discrete information relating to recent displacement reversals, or *endpoints*.

The topic of memory brings us finally to a common point across both directions: many of these models show some form of memory, *i*.*e*., a dependency on past history. The Biot element shows continuous memory, as per its memory function *Q*_BI_ (Fig. 2A); whereas hysteretic models typically show discrete (endpoint) memory. Both memory effects would be testable with current exoskeletal DMA techniques, *cf*. Guyer et al. (1997) and Wold et al. (2023). Structural memory is an area which is both understudied and has the potential to provide significant insight into the nature and origins of exoskeletal structural damping.

### 6.3. Conclusion

In this work, we developed the first complete time-domain models of rate- (or, frequency-) independent damping in insect exoskeleta. We began by translating an established frequency-domain exoskeletal damping model, the *iγ* model, into the time domain; identifying this time-domain representation as the well-known singular integral operator, the Hilbert transform. This process reveals that this established model is strongly non-causal, *i*.*e*., it violates the directionality of time. While this noncausality is not particularly problematic under approximately sinusoidal motion, it is more serious under isolated perturbations—for these, significant anticipatory force generation occurs. In response, we extend results from seismic analysis to develop *causal* time-domain models of linear rate-independent damping in the exoskeleton, including an extended form of the Biot element. Recognising these time-domain models as strongly singular integrals defined by Hadamard finite-part integration, we develop and validate numerical quadrature techniques for them—allowing us to compute responses to impulses, perturbations, and general motion; and simulate the damper within wider structural models. We apply these techniques to several problems in insect biomechanics: identifying thoracic damping from non-sinusoidal DMA data in the presence of nonlinear elasticity; identifying complete limb structural parameters from perturbation kinematics; and simulating exoskeletal structural models in time. However, in doing so, we identify a key caveat with both the *iγ* damping model and its causal analogues: they are not purely dissipative, but are capable of generating negative power, and in ways that are inconsistent with energy storage from a single nonlinear elasticity. Together, these methods provide (**i**) rigorous and complete methods to identify and simulate rate-independent damping in various exoskeletal contexts; (**ii**) testable predictions to distinguish different models of, and mechanisms behind, exoskeletal damping; and (**iii**) avenues for further improvement in modelling and understanding this understudied property of the insect exoskeleton.

## Data availability

Code and data will be made available upon publication.

## Footnotes

† Hadamard’s finite part is not the only option for regularisation, *cf*. Gel’fand and Shilov (1964) and Lighthill (1958). As per Estrada and Kanwal (1989), different regularisations can differ by up to a factor of *δ*, which be a source of confusion—for instance, seismic analysis literature (Luo and Ikago, 2021; Makris, 1997a), following Lighthill (1958), selects regularisations which eliminate the *δ* term in Eq. 14. Unfortunately, the differing factor of *δ* is crucial, as it determines an effective stiffness term, *Kx*(*t*), in the damper’s response (Eq. 15). When we turn to numerical computation in §3, we will be able to partially resolve this ambiguity via a numerical method for estimating the appropriate *δ* factor.

‡ Does conservation of energy (in time) even have any meaning within a non-causal (time-violating) structure? A deeper theoretical study would be required to resolve this.

