## Supplementary Material for "Modelling rate-independent damping in insect exoskeleta via singular integral operators"

### S1. Memory function of the Biot element

To show that Eq. 19 of the main text is the memory function of the Biot element, note that we may express the Fourier transform of the key distribution within $Q_{\mathrm{BI}}(t)$, $\mathrm{f.p.}\,\Theta(t)e^{-\Omega_{\mathrm{trans}}|t|}/t$, via the Fourier convolution theorem as:

$$\mathcal{F}\left\{\mathrm{f.p.}\frac{\Theta(t)}{t}e^{-\Omega_{\mathrm{trans}}|t|}\right\}=\frac{1}{2\pi}\,\mathcal{F}\left\{\mathrm{p.v.}\frac{e^{-\Omega_{\mathrm{trans}}|t|}}{t}\right\}*\mathcal{F}\{\Theta(t)\}+D, \tag{1.1}$$

where we introduce the indeterminate factor $D\delta$ to account for the fact that switching regularisation methods may cause us to lose a delta distribution in the time domain (Estrada and Kanwal, 1989), *i.e.*, a constant in the frequency domain. We will compute the value of $D$ later. Then, note the established results for the individual Fourier transforms within the convolution on the RHS of Eq. 1.1:

$$\begin{aligned}\mathcal{F}\left\{\mathrm{p.v.}\frac{e^{-\Omega_{\mathrm{trans}}|t|}}{t}\right\}&=-2i\tan^{-1}\left(\frac{\Omega}{\Omega_{\mathrm{trans}}}\right),\\ \mathcal{F}\{\Theta(t)\}&=-i\mathrm{p.v.}\frac{1}{\Omega}+\pi\delta(\Omega),\end{aligned} \tag{1.2}$$

derivable either via hyperfunctions (Graf, 2010) or more classical approaches (Badrieh, 2018; Oberhettinger, 1990). Substituting these terms, the right-hand side of Eq. 1.1 reduces to the computation of a Hilbert transform (the convolution of a function with $\mathrm{p.v.}\,1/\Omega$, *i.e.* $\mathcal{H}$, see Eq. 5 of the main text):

$$\mathcal{F}\left\{\mathrm{f.p.}\frac{\Theta(t)}{t}e^{-\Omega_{\mathrm{trans}}|t|}\right\}=-\mathcal{H}\left\{\tan^{-1}\left(\frac{\Omega}{\Omega_{\mathrm{trans}}}\right)\right\}-i\tan^{-1}\left(\frac{\Omega}{\Omega_{\mathrm{trans}}}\right)+D\,. \tag{1.3}$$

Using further established results for the Hilbert transform (King, 2009), this evaluates to:

$$\mathcal{F}\left\{\mathrm{f.p.}\frac{\Theta(t)}{t}e^{-\Omega_{\mathrm{trans}}|t|}\right\}=-\ln\sqrt{1+\left(\frac{\Omega}{\Omega_{\mathrm{trans}}}\right)^2}-i\tan^{-1}\left(\frac{\Omega}{\Omega_{\mathrm{trans}}}\right)+D. \tag{1.4}$$

Eq. 1.4 is the complex modulus of the Biot element, $\widehat{\mathbb{E}}_{\mathrm{BI}}(\Omega)$ (Eq. 16 of the main text), up to a factor of $-2\eta/\pi$, the term in $K_{\mathrm{BI}}$, and for some term $D$.

Computing $D$ is not necessary to make numerical use of the model (see §3.2 of the main text), but the term is useful to have. To compute $D$ we use an *ad hoc* method based on the response of the model at zero frequency ($\Omega=0$). At $\Omega=0$:

$$\widehat{\mathbb{E}}_{\mathrm{BI}}(0)=-\frac{2\eta}{\pi}K_{\mathrm{BI}}, \tag{1.5}$$

that is, the element behaves as a linear spring with stiffness $\widehat{\mathbb{E}}_{\mathrm{BI}}(0)$. Then consider a time-domain evaluation of the damper's response to a constant input, $x = 1$. By Eq. 1.4, the memory function of the element is:

$$Q_{\mathrm{BI}}(t) = -\frac{2\eta}{\pi}\,\mathrm{f.p.}\frac{\Theta(t)}{t}e^{-\Omega_{\mathrm{trans}}|t|} - \frac{2\eta}{\pi}K_{\mathrm{BI}}\delta(t) + C\delta(t), \tag{1.6}$$

with unknown factor $C$. The constant response, $E_0$, of Eq. 1.6 to $x = 1$ should equal $\widehat{\mathbb{E}}_{\mathrm{BI}}(0)$, and we seek to compute $C$ such that this occurs. $E_0$ is given by:

$$\begin{aligned} E_0 &= \left(-\frac{2\eta}{\pi}\,\mathrm{f.p.}\frac{\Theta(t)}{t}e^{-\Omega_{\mathrm{trans}}|t|} - \frac{2\eta}{\pi}K_{\mathrm{BI}}\delta(t) + C\delta(t)\right) * 1 \\ &= -\frac{2\eta}{\pi}\mathrm{f.p.}\int_{-\infty}^{\infty}\frac{\Theta(\tau)}{\tau}e^{-\Omega_{\mathrm{trans}}|\tau|}\,d\tau - \frac{2\eta}{\pi}K_{\mathrm{BI}} + C \\ &= -\frac{2\eta}{\pi}\mathrm{f.p.}\int_{0}^{\infty}\frac{e^{-\Omega_{\mathrm{trans}}\tau}}{\tau}\,d\tau - \frac{2\eta}{\pi}K_{\mathrm{BI}} + C. \end{aligned} \tag{1.7}$$

With a change of variables, $\nu = \Omega_{\mathrm{trans}}\tau$ in the finite-part integral, we can identify its value with the Hadamard regularisation of the Gamma function ($\Gamma$) (Paycha, 2012), which for consistency We denote $\mathrm{f.p.}\,\Gamma$ *in lieu* of Paycha's (2012) $\Gamma^{\mathrm{Had}}$:

$$\begin{aligned} E_0 &= -\frac{2\eta}{\pi}\mathrm{f.p.}\int_{0}^{\infty}\frac{e^{-\nu}}{\nu}\,d\nu - \frac{2\eta}{\pi}K_{\mathrm{BI}} + C. \\ &= -\frac{2\eta}{\pi}\mathrm{f.p.}\,\Gamma(0) - \frac{2\eta}{\pi}K_{\mathrm{BI}} + C. \end{aligned} \tag{1.8}$$

$\mathrm{f.p.}\,\Gamma(0) = -\gamma_e$, the Euler-Mascheroni constant. So, for consistency:

$$\begin{aligned} &E_0 = \widehat{\mathbb{E}}_{\mathrm{BI}}(0) \\ &\therefore \frac{2\eta}{\pi}\gamma_e - \frac{2\eta}{\pi}K_{\mathrm{BI}} + C = -\frac{2\eta}{\pi}K_{\mathrm{BI}}\,, \\ &\therefore C = -\frac{2\eta}{\pi}\gamma_e. \end{aligned} \tag{1.9}$$

This constant is identical to that that appears in the causal hysteretic damper (Eq. 14 of the main text). We may express:

$$Q_{\mathrm{BI}}(t) = -\frac{2\eta}{\pi}\,\mathrm{f.p.}\frac{\Theta(t)}{t}e^{-\Omega_{\mathrm{trans}}|t|} - \frac{2\eta}{\pi}(\gamma_e + K_{\mathrm{BI}})\delta(t), \tag{1.10}$$

as per Eq. 19 of the main text.

### S2. Time-domain stencils for finite-part integrals

#### S2.1. Overview of finite-part integration

In this section we provide a brief overview of finite-part integration, and a detailed derivation of the time-domain numerical method for computing the finite-part integrals within the causal rate-independent damper and the Biot element. *First*, regarding finite-part integration—here, we follow Estrada and Kanwal (1994, §2.4), Jones (1996) and Kanwal (2004, §4.2); but additional details can be found in Gel'fand and Shilov (1964). The central concept of the Hadamard finite part is that many types of divergent integrals (*i.e.*, nominally of infinite value) can be split into a diverging term and a unique finite converging term. By discarding the diverging term, the 'finite part' is recovered. Note that this finite part integral *no longer* represents the 'area under the curve' in an intuitive sense: for instance, a finite-part integral may be negative-valued when the integrand is strictly positive. Such an integral might appear to be meaningless, but finite-part integrals of this type arise in certain analyses of real physical phenomena. They have arisen here, in the analysis of rate-independent damping models, and so a method to compute them is required.

To define finite part integration quantitively, consider first the general integral:

$$\int_0^L \frac{\phi(\tau)}{\tau} d\tau \, , \tag{2.1}$$

where $\phi(\tau)$ is some function that is integrable over $[0, L]$ (*i.e.*, its own absolute integral converges to a finite value). However, because of the factor $1/\tau$, Eq. 2.1 is a diverging integral, and the point of divergence (singularity) is $\tau = 0$. To see this divergence in action, consider the related integral

$$\int_{\epsilon_1}^L \frac{\phi(\tau)}{\tau} d\tau \, , \tag{2.2}$$

for some small $\epsilon_1$. We see that Eq. 2.2 has finite value for finite $\epsilon_1$, but this value does not converge as $\epsilon_1 \to 0$; instead, the integral $\to \pm\infty$. To separate the converging and diverging terms within the integral, we perform the following process. We split the integral up by adding another arbitrary reference point $\epsilon_2$: integration is split into the intervals $[\epsilon_1, \epsilon_2]$ and $[\epsilon_2, L]$. We then perform what appears as a mathematical trick, by adding the term $\phi(0) - \phi(0) = 0$, yielding:

$$\begin{aligned}
\int_{\epsilon_1}^L \frac{\phi(\tau)}{\tau} d\tau &= \int_{\epsilon_1}^{\epsilon_2} \frac{\phi(\tau)}{\tau} d\tau + \int_{\epsilon_2}^L \frac{\phi(\tau)}{\tau} d\tau \\
&= \int_0^{\epsilon_2} \frac{\phi(\tau) + \phi(0) - \phi(0)}{\tau} d\tau + \int_{\epsilon_2}^L \frac{\phi(\tau)}{\tau} d\tau \\
&= \int_{\epsilon_1}^{\epsilon_2} \frac{\phi(0)}{\tau} d\tau + \int_{\epsilon_1}^{\epsilon_2} \frac{\phi(\tau) - \phi(0)}{\tau} d\tau + \int_{\epsilon_2}^L \frac{\phi(\tau)}{\tau} d\tau \\
&= \underbrace{\phi(0) \ln \epsilon_2}_{A} - \underbrace{\phi(0) \ln \epsilon_1}_{B} + \underbrace{\int_{\epsilon_1}^{\epsilon_2} \frac{\phi(\tau) - \phi(0)}{\tau} d\tau}_{C} + \underbrace{\int_{\epsilon_2}^L \frac{\phi(\tau)}{\tau} d\tau}_{D} \, ,
\end{aligned} \tag{2.3}$$

Now let $\epsilon_2 \to 0$. The terms $A$, $C$ and $D$ converge when taking this limit (Estrada and Kanwal, 1994). Term $B$ diverges: to define a finite value for the integral in Eq. 2.3, we thus discard this term. This allows us to definite the finite part integral as:

$$\text{f.p.}\int_0^L \frac{\phi(\tau)}{\tau}d\tau = \phi(0)\ln\epsilon + \int_0^\epsilon \frac{\phi(\tau)-\phi(0)}{\tau}d\tau + \int_\epsilon^L \frac{\phi(\tau)}{\tau}d\tau \qquad (2.4)$$

where we now denote $\epsilon = \epsilon_2$. Crucially, this $\epsilon$ was a free parameter, and we can confirm (Estrada and Kanwal, 1989; 1994) that Eq. 2.4 gives the same (finite!) result independent of $\epsilon$—we are free to choose, based on numerical or analytical convenience. It is thus typical in this distributional literature to choose $\epsilon = 1$, because then $\ln\epsilon = 0$ and Eq. 2.4 simplifies.

**S2.2. The causal rate-independent damper: numerical method with analytical corrector**

Eq. 2.4 readily yields a basic numerical method for computing the responses of the causal rate-independent damper in the time domain. The time-domain formulation of this damper's response is:

$$\begin{aligned} F_{\text{CRI}}(t) &= -\frac{2\eta}{\pi}\text{f.p.}\int_{-\infty}^t \frac{x(\tau)}{t-\tau}d\tau - \frac{2\eta}{\pi}(\gamma_e + \ln|\Omega_{\text{ref}}|)x(t), \\ &= -\frac{2\eta}{\pi}\text{f.p.}\int_0^\infty \frac{x(t-\tau)}{\tau}d\tau - \frac{2\eta}{\pi}(\gamma_e + \ln|\Omega_{\text{ref}}|)x(t), \end{aligned} \qquad (2.5)$$

within which is the finite-part integral:

$$\text{f.p.}\int_0^\infty \frac{x(t-\tau)}{\tau}d\tau\,. \qquad (2.6)$$

At this point, to be specific, we restrict ourselves to cases where $x(t-\tau)$ is a function of compact support, *cf.* Kanwal (1997) and Nussenzveig (1972), *i.e.*, input data of finite length. This condition that always be satisfied by physical time-series input data. In this case we can approximate:

$$\text{f.p.}\int_0^\infty \frac{x(t-\tau)}{\tau}d\tau \cong \text{f.p.}\int_0^L \frac{x(t-\tau)}{\tau}d\tau\,, \qquad (2.7)$$

for some $L$, and we know that for sufficiently large $L$ this approximation will be exact. With that formality established, we can identify Eq. 2.7 with Eq. 2.4 under $\phi(\tau) = x(t-\tau)$, and so:

$$\begin{aligned} &\text{f.p.}\int_0^L \frac{x(t-\tau)}{\tau}d\tau \\ &\qquad = x(t)\ln\epsilon + \int_0^\epsilon \frac{x(t-\tau)-x(t)}{\tau}d\tau + \int_\epsilon^L \frac{x(t-\tau)}{\tau}d\tau\,. \end{aligned} \qquad (2.8)$$

Then consider the nature of the function $x(t-\tau)$. For numerical input data, this function is not itself continuous but discrete: its values are given at discrete points (*e.g.*, defined in an array $\mathbf{x}$, with elements $x_i$, over some discrete time array $\boldsymbol{\tau}$). Here we assume that $\boldsymbol{\tau}$ takes the form $\boldsymbol{\tau} = [0, \Delta\tau, \ldots, L]$, *i.e.*, only the first timestep $\Delta\tau$ is relevant. Over this first interval, $[0, \Delta\tau]$, we make the zeroth-order approximation, that $x(\tau) = x(0)$, that is, the continuous value is constant over this timestep. If we make this approximation in Eq. 2.8, and take $\epsilon = \Delta\tau$, then the second term vanishes and we have:

$$\text{f.p.}\int_0^L \frac{x(t-\tau)}{\tau}d\tau \cong x(t)\ln\Delta t + \int_{\Delta t}^L \frac{x(t-\tau)}{\tau}d\tau\,. \qquad (2.9)$$

The remaining integral in Eq. 2.9, over $[\Delta\tau, L]$ is an ordinary integral that can be approximated by any relevant quadrature method (trapezoidal, Simpson's, Gaussian). In this article's implementation, Simpson's 1/3 rule is used, with a corrector for the case of an even number of intervals. Returning to the force response of the damper, we have the numerical approximation:

$$\begin{aligned} F_{\mathrm{CRI}}(t) &= -\frac{2\eta}{\pi}\,\mathrm{f.p.}\int_0^\infty \frac{x(t-\tau)}{\tau}\,d\tau - \frac{2\eta}{\pi}(\gamma_e + \ln|\Omega_{\mathrm{ref}}|)x(t), \\ &\cong -\frac{2\eta}{\pi}\underbrace{\int_{\Delta t}^{L} \frac{x(t-\tau)}{\tau}\,d\tau}_{\substack{\text{conventional}\\ \text{quadrature}}} - \frac{2\eta}{\pi}(\gamma_e + \ln|\Omega_{\mathrm{ref}}| + \ln\Delta t)x(t), \end{aligned} \tag{2.10}$$

**S2.3. The Biot element: numerical method with analytical corrector**

We can analyse the Biot element along similar lines. The time-domain formulation of the Biot element's response is:

$$\begin{aligned} F_{\mathrm{BI}}(t) &= -\frac{2\eta}{\pi}\,\mathrm{f.p.}\int_{-\infty}^{t} \frac{x(\tau)}{t-\tau} e^{-\Omega_{\mathrm{trans}}|\tau-t|}\,d\tau - \frac{2\eta}{\pi}(\gamma_e + K_{\mathrm{BI}})x(t), \\ &= -\frac{2\eta}{\pi}\,\mathrm{f.p.}\int_0^\infty \frac{x(t-\tau)}{\tau} e^{-\Omega_{\mathrm{trans}}\tau}\,d\tau - \frac{2\eta}{\pi}(\gamma_e + K_{\mathrm{BI}})x(t), \end{aligned} \tag{2.11}$$

within which is the finite-part integral:

$$\mathrm{f.p.}\int_0^\infty \frac{x(t-\tau)}{\tau} e^{-\Omega_{\mathrm{trans}}\tau}\,d\tau \cong \mathrm{f.p.}\int_0^L \frac{x(t-\tau)}{\tau} e^{-\Omega_{\mathrm{trans}}\tau}\,d\tau\,. \tag{2.12}$$

Considering again discrete $x$ ($\mathbf{x}$) over time array $\boldsymbol{\tau} = [0, \Delta\tau, \dots, L]$, we make the same zeroth-order approximation, $x(\tau) = x(0)$, and so can split Eq. 2.6 directly into:

$$\begin{aligned} &\mathrm{f.p.}\int_0^L \frac{x(t-\tau)}{\tau} e^{-\Omega_{\mathrm{trans}}\tau}\,d\tau \\ &\quad = x(t)\,\mathrm{f.p.}\int_0^{\Delta\tau} \frac{e^{-\Omega_{\mathrm{trans}}\tau}}{\tau}\,d\tau + \int_{\Delta\tau}^{L} \frac{x(t-\tau)}{\tau} e^{-\Omega_{\mathrm{trans}}\tau}\,d\tau\,, \end{aligned} \tag{2.13}$$

where the second integral on the RHS is now again an ordinary integral. We can rearrange the remaining finite-part integral into an expression involving the exponential integral, the special function styled Ei(·):

$$\begin{aligned} \mathrm{f.p.}\int_0^{\Delta\tau} \frac{e^{-\Omega_{\mathrm{trans}}\tau}}{\tau}\,d\tau &= \mathrm{f.p.}\int_0^{\Omega_{\mathrm{trans}}\Delta\tau} \frac{e^{-v}}{v}\,dv\,, \\ &= \mathrm{f.p.}\int_0^\infty \frac{e^{-v}}{v}\,dv - \mathrm{p.v.}\int_{\Omega_{\mathrm{trans}}\Delta\tau}^{\infty} \frac{e^{-v}}{v}\,dv\,, \\ &= \mathrm{f.p.}\,\mathrm{Ei}(0) + \mathrm{Ei}(-\Omega_{\mathrm{trans}}\Delta\tau)\,, \end{aligned} \tag{2.14}$$

where

$$\mathrm{Ei}(y) = -\int_{-y}^{\infty} \frac{e^{-v}}{v}\,dv\,. \tag{2.15}$$

and $\mathrm{Ei}(0)$ must be defined by finite part. From Paycha (2012), $\text{f.p.}\,\mathrm{Ei}(0) = -\gamma_e$. As $\mathrm{Ei}(\Omega_{\text{trans}}\Delta\tau)$ can be evaluated using standard methods (MATLAB function `expint`), we now have a complete numerical method for computing the Biot element's response. We have the finite-part evaluation:

$$\begin{aligned} &\text{f.p.} \int_0^L \frac{x(t-\tau)}{\tau} e^{-\Omega_{\text{trans}}\tau} d\tau \\ &\qquad = (-\gamma_e + \mathrm{Ei}(-\Omega_{\text{trans}}\Delta\tau))x(t) \; + \int_{\Delta\tau}^L \frac{x(t-\tau)}{\tau} e^{-\Omega_{\text{trans}}\tau} d\tau\,, \end{aligned} \tag{2.16}$$

which, when substituted into Eq. 2.12 and 2.11, leads to cancellation of the $\gamma_e$ term, and thus:

$$\begin{aligned} F_{\text{BI}}(t) &= -\frac{2\eta}{\pi}\,\text{f.p.} \int_0^\infty \frac{x(t-\tau)}{\tau} e^{-\Omega_{\text{trans}}\tau} d\tau - \frac{2\eta}{\pi}(\gamma_e + K_{\text{BI}})x(t), \\ &\cong -\frac{2\eta}{\pi} \underbrace{\int_{\Delta t}^L \frac{x(t-\tau)}{\tau} e^{-\Omega_{\text{trans}}\tau} d\tau}_{\substack{\text{conventional} \\ \text{quadrature}}} - \frac{2\eta}{\pi}(K_{\text{BI}} + \mathrm{Ei}(-\Omega_{\text{trans}}\Delta t))x(t), \end{aligned} \tag{2.17}$$

**S2.4. Numerical correctors for the finite part**

In Eq. 2.10 and 2.17, the coefficients of $x(t)$, *i.e.*, the factors of $\gamma_e + \ln|\Omega_{\text{ref}}| + \ln\Delta t$ and $K_{\text{BI}} + \mathrm{Ei}(-\Omega_{\text{trans}}\Delta t)$ are effective stiffnesses that correct the ordinary integral term into the relevant finite-part response, tuned to the appropriate reference frequency ($\Omega_{\text{ref}}$). As we have seen, significant analytical effort is required to derive these corrector terms. However, we observe that there is an alternate method to compute them, which both validates the analytical correctors and provides an accurate and general technique to potentially replace them.

Considering the causal rate-independent damper as example, we observe the analytical results, Eq. 2.10, indicate that the ordinary integral over $[\Delta t, L]$ can be corrected to the true finite part over $[0, L]$ via some effective stiffness which we term $C_{\text{CRI}}$:

$$F_{\text{CRI}}(t) \cong -\frac{2\eta}{\pi}\int_{\Delta t}^L \frac{x(t-\tau)}{\tau} d\tau - C_{\text{CRI}}x(t). \tag{2.18}$$

A defining feature of this correction, and thus $C_{\text{CRI}}$, is that the storage modulus of the damper at the frequency $\Omega_{\text{ref}}$ should be zero. We can estimate this storage modulus directly using Eq. 2.18 by testing the response to a sinusoid $x(t) = \cos(\Omega_{\text{ref}}t)$ and observing the force at $t = 0$:

$$E'_{\text{CRI}}(\Omega_{\text{ref}}) \cong -\frac{2\eta}{\pi}\int_{\Delta t}^L \frac{\cos(-\Omega_{\text{ref}}\tau)}{\tau} d\tau - C_{\text{CRI}}. \tag{2.19}$$

Then, solving for $E'_{\text{CRI}}(\Omega_{\text{ref}}) = 0$, we obtain:

$$C_{\text{CRI}} \cong -\frac{2\eta}{\pi}\int_{\Delta t}^L \frac{\cos(\Omega_{\text{ref}}\tau)}{\tau} d\tau\,, \tag{2.20}$$

which is a numerical estimate for $C_{\text{BI}}$ which can (and should) be performed by the same quadrature method as used to compute the ordinary damper response (Eq. 2.10). The reason for selecting the same quadrature is that, if there are any errors in the quadrature that specifically lead to an effective stiffness term, this method will eliminate them.

An identical principle can be applied to the Biot element, with only the integral kernel changing. We have:

$$F_{\mathrm{BI}}(t) \cong -\frac{2\eta}{\pi}\int_{\Delta t}^{L}\frac{x(t-\tau)}{\tau}d\tau - C_{\mathrm{BI}}x(t), \tag{2.21}$$

with:

$$C_{\mathrm{BI}} \cong -\frac{2\eta}{\pi}\int_{\Delta t}^{L}\frac{\cos(\Omega_{\mathrm{ref}}\tau)}{\tau}e^{-\Omega_{\mathrm{trans}}\tau}d\tau\,, \tag{2.22}$$

and the relationship between $\Omega_{\mathrm{ref}}$ and $\Omega_{\mathrm{trans}}$ being determined by the selected $N_{\mathrm{ref}}$. These two numerical correctors, Eq. 2.20 and Eq. 2.22, are Eq. 24 of the main text.
